# Single-cell dissection of the human cerebrovasculature in health and disease

**DOI:** 10.1101/2021.04.26.440975

**Authors:** Francisco J. Garcia, Na Sun, Hyeseung Lee, Brianna Godlewski, Kyriaki Galani, Julio Mantero, David A. Bennett, Mustafa Sahin, Manolis Kellis, Myriam Heiman

## Abstract

Despite the importance of the blood-brain barrier in maintaining normal brain physiology and in understanding neurodegeneration and CNS drug delivery, human cerebrovascular cells remain poorly characterized due to their sparsity and dispersion. Here, we perform the first single-cell characterization of the human cerebrovasculature using both *ex vivo* fresh-tissue experimental enrichment and *post mortem in silico* sorting of human cortical tissue samples. We capture 31,812 cerebrovascular cells across 17 subtypes, including three distinct subtypes of perivascular fibroblasts as well as vasculature-coupled neurons and glia. We uncover human-specific expression patterns along the arteriovenous axis and determine previously uncharacterized cell type-specific markers. We use our newly discovered human-specific signatures to study changes in 3,945 cerebrovascular cells of Huntington’s disease patients, which reveal an activation of innate immune signaling in vascular and vasculature-coupled cell types and the concomitant reduction to proteins critical for maintenance of BBB integrity. Finally, our study provides a comprehensive resource molecular atlas of the human cerebrovasculature to guide future biological and therapeutic studies.

## Introduction

The cerebrovasculature exhibits specialized barrier properties that regulate the transport of biomolecules and maintain brain homeostasis^1,2^. Structural imaging and molecular studies have yielded important insights into the mouse cerebrovasculature^3,4^. However, human brains exhibit increased complexity and energetic needs, likely accompanied by human-specific adaptations of the cerebrovasculature, which remain uncharacterized^1^. Moreover, cerebrovascular dysfunction and blood-brain barrier (BBB) breakdown are hypothesized to play important roles in aging^5^ and in multiple brain disorders^1,6^, including Huntington’s disease (HD)^7,8^, Alzheimer’s disease (AD)^9^, and cerebrovascular diseases such as vascular dementia (e.g. CADASIL)^10^, primary familial brain calcification (PFBC)^11^, and cerebral amyloid angiopathy (CAA)^12^. Thus, understanding the human cerebrovasculature and BBB, both in physiologic conditions and in pathological conditions that break down their integrity, is a pressing need for both scientific and clinical reasons.

Here, we address this challenge by reporting the first comprehensive single-cell molecular atlas of human cerebrovasculature cells, across ~32,000 cells from both (a) *ex vivo* freshly-resected surgical human brain tissue coupled with a new cerebrovascular cell enrichment protocol, and (b) an *in silico* cell sorting method from *post mortem* frozen human brain tissue. We computationally integrate these datasets to characterize 17 subtypes of cells in the human BBB, including three perivascular fibroblast cell subtypes and a subset of vasculature-coupled brain cells. Our studies reveal unique human-specific transcriptomic signatures along the arteriovenous axis for endothelial and mural cells and elucidate dysregulated transcriptional changes of cell types comprising the human cerebrovasculature in HD. Our study highlights the unique cell type-specific and species-specific characteristics in humans and provides a framework for mechanistic dissection of pathological state dysfunctions.

## Results

### Fresh-tissue cerebrovasculature cell enrichment

Due to their low abundance in the brain and poor capture efficiency with droplet-based sequencing methodologies, endothelial and mural cells of the cerebrovasculature have been challenging to characterize at single-cell resolution. Prior mouse molecular profiling studies have used transgenic reporter lines and fluorescence-activated cell sorting (FACS) to enrich for cerebrovasculature cells^3,13^, but these methods are incompatible with human tissue studies. To address this challenge, we developed a Blood Vessel Enrichment (BVE) protocol (**Fig. 1a**, top) for enriching human vascular cell types from fresh and fresh-frozen brain tissue for single-cell applications by homogenizing tissue samples, performing dextran-based density ultracentrifugation^14^, and using the microvessel-enriched pellet as input for single-nucleus RNA-seq profiling^15^. We validated our protocol using mouse cortical samples, showing enrichment for markers of endothelial (*Abcb1a, Cldn5, Mfsd2a*) and mural (*Pdgfrb, Acta2, Myh11*) cells (**Extended Data Fig. 1a**), but no enrichment for markers of neurons (*Rbfox3*), oligodendrocytes (*Mog*), microglia (*Aif1*), or somatic astrocytes (*Aldh1l1*). However, we found that our protocol still captures: i) astrocytic end-feet (marked by *Aqp4*)^*14*^, as they remain attached to the vasculature post-enrichment, which we confirmed by indirect immunofluorescence (**Extended Data Fig. 1b**), and also ii) vasculature-coupled glial and neuronal cells, as we discuss below.

**Figure 1.**
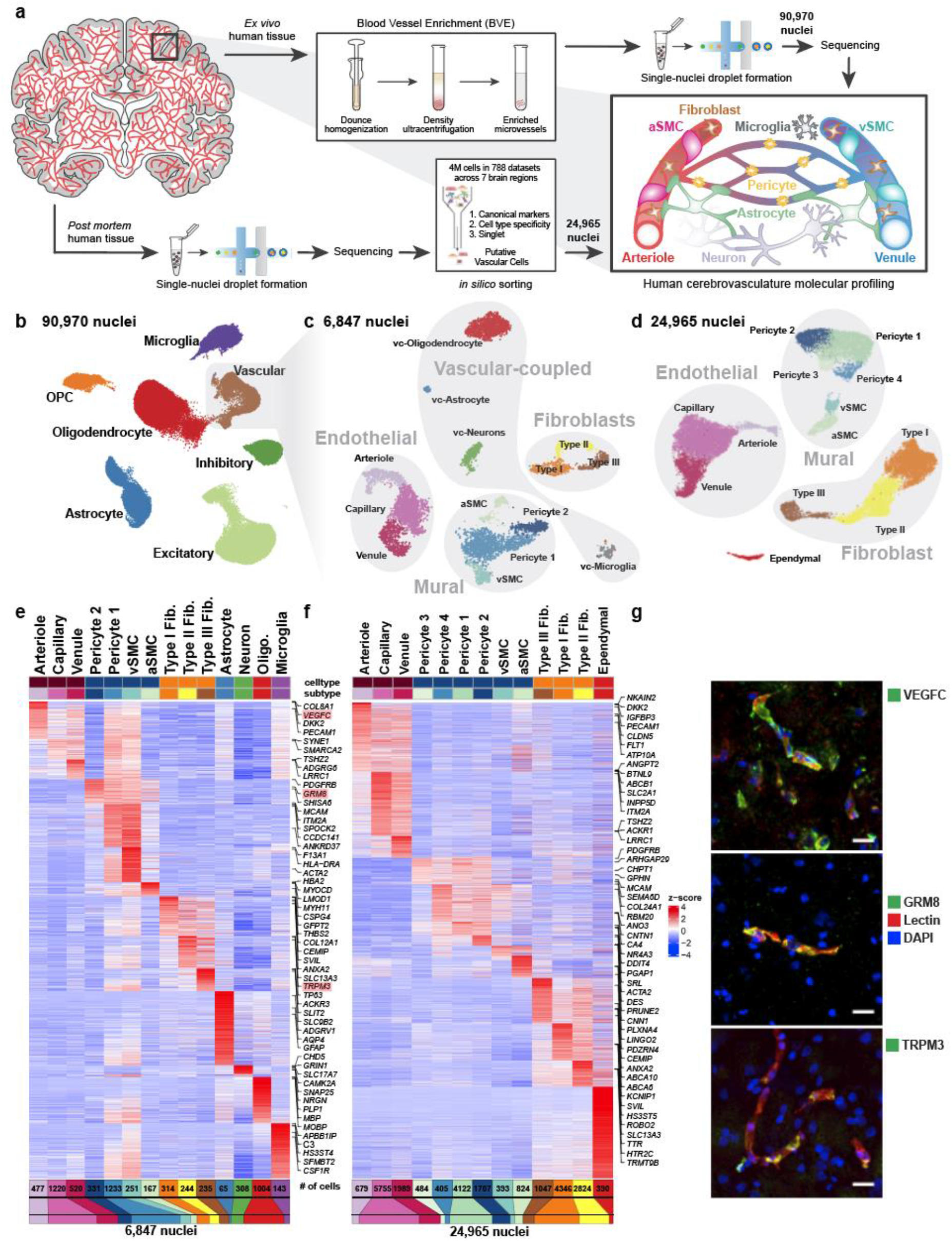
snRNA-seq profiling of the human cerebrovasculature. **a**. Experimental schematic. **b**. Global Uniform Manifold Approximation and Projection (UMAP) of 90,970 profiled nuclei from *ex vivo* human temporal cortex. **c**. UMAP of 6,847 profiled vascular nuclei from the highlighted cluster in **b. d**. UMAP of 24,965 *in silico* sorted vascular nuclei from *post mortem* human brains. **e,f**. Highly expressed genes of sub-clusters in **c** and **d**, respectively. **g**. Validation of markers *VEGFC, GRM8*, and *TRPM3* by indirect immunofluorescence staining (each in green pseudocolor). Scale bar, 20μm.

We applied our BVE protocol to the study of *ex vivo* fresh-frozen tissue from 17 live temporal lobe surgical resections of patients with intractable epilepsy (**Supplementary Table 1**), thus providing samples free from hypoxia and other death-induced environmental stressors and avoiding decreased RNA quality associated with long *post mortem* intervals^16^. We specifically selected tissue distal to the epileptic focus, as has been recently performed for electrophysiological characterization of human neurons^17^. For seven samples, we used standard snRNA-seq^18^ without the BVE protocol to confirm the absence of BVE-induced biases (**Extended Data Fig. 2a,b**). We obtained 90,970 single nuclei after quality control (**Fig. 1b, Extended Data Fig. 2c-e**), including 16 subtypes of excitatory neurons, 6 subtypes of inhibitory neurons^19^, oligodendrocytes, oligodendrocyte precursor cells, astrocytes, microglia, and 6,847 vascular cells, a 20-fold enrichment from previous studies^15^ (8% vs. ~0.4%). We distinguished 14 subtypes of vascular cells (**Fig. 1c**), including: (a) 3 subtypes of endothelial cells (*n*=2,552), separated along the arteriovenous axis; (b) 4 subtypes of mural cells (*n*=1,982), including 1,564 pericytes and 418 smooth muscle cells (SMCs), showing a pericyte-SMC gradation; (c) 3 subtypes of perivascular fibroblasts (*n*=793) with distinct marker genes; and (d) 4 subtypes of vasculature-coupled (vc) neuronal and glial cells (*n*=1,520), consisting of vc-neurons, vc-oligodendrocytes, vc-astrocytes, and vc-microglia.

### *Post mortem* tissue *in silico* cerebrovasculature cell sorting

We complemented our BVE-captured cerebrovasculature cell compendium with *in silico* sorting of 24,965 cerebrovascular cells from *post mortem* samples across 7 different brain regions, including prefrontal cortex, mid-temporal cortex, angular gyrus, entorhinal cortex, thalamus, hippocampus and mammillary body, from the ROSMAP studies^20^. We used a combination of canonical vasculature markers and whole-transcriptome cellular signatures, separately from our BVE protocol, to ensure independence (**Fig. 1a**, bottom, **Methods**). Using canonical markers, gene-set enrichments, and mouse cerebrovasculature data^3^, we annotated 13 subtypes of vascular cells (**Fig. 1d**), including: (a) 3 subtypes of endothelial cells (*n*=8,423) separated along the arteriovenous axis; (b) 6 subtypes of mural cells (*n*=7,935), including 6,718 pericytes, separated into 4 subtypes and 1,217 smooth muscle cells (SMCs), also showing a pericyte-SMC gradation; (c) 3 subtypes of fibroblasts (*n*=8,217), with matching subtypes to *ex vivo*; and (d) an ependymal cell subcluster (*n*=390).

Remarkably, the majority of cerebrovascular subtypes in the *ex vivo* BVE dataset were also found in our *in silico post mortem* sorted dataset, validating the consistency of cell types across experimental platforms and sample cohorts. Pericytes exhibited greater heterogeneity in the *in silico* sorted dataset (4 subtypes vs. 2 subtypes) though this may be due to a larger number of profiled cells (6,718 vs. 1,564) compared to the *ex vivo* dataset. Notably, the subset of vasculature-coupled canonical cell types (vc-neurons, vc-oligodendrocytes, vc-astrocytes, and vc-microglia) was only found in the *ex vivo* dataset, while a small cluster of ependymal cells was specific to the *in silico* sorted data. As vasculature-coupled cell types are co-enriched by our BVE protocol, their presence in the *ex vivo* dataset and not the total (ROSMAP) snRNA-seq preparation from *post mortem* tissue is expected. Similarly, as ependymal cells are the constituents of ventricles producing cerebrospinal fluid, it is not expected that they would be present in the very precisely resected *ex vivo* surgical tissue samples.

### Genes and pathways defining each cell type

We next used our snRNA-seq data to create an atlas of human cerebrovascular cells and reveal their defining molecular characteristics by combining our two independent experimental and computational enrichment strategies. We found robust expression a total of 17,212 genes (each in 50+ cells), including 4,090 differentially expressed genes (DEGs) among cell types in the *ex vivo* and 3,997 DEGs in *post mortem* datasets, which vastly expand the previously-known set of canonical markers for each cell type, and enable us to infer candidate functional roles for each cell type. We confirmed robust differential expression for known marker genes in endothelial (*CLDN5*), mural (*TAGLN*), fibroblast (*CEMIP*), ependymal (*TTR*), astroglia (*GFAP*), neurons (*CAMK2A*), oligodendrocytes (*PLP1*), and microglial (*CSF1R*) cell types (**Fig. 1e-f, Supplementary Table 2**). In addition, we identified markers for subtypes of these cells, separating endothelial cells into arteriole (*VEGFC, ARL15*), capillary (*MFSD2A, SLC7A5*), venule (*TSHZ2, ADGRG6*), separating mural cells into aSMC (*ACTA2, MYH11*), vSMC (*MRC1, CD74*), pericytes (*GRM8, PDGFRB*), and separating fibroblasts into Type I (*ABCA10, FBLN1*), Type II (*TRPM3, MYRIP*), and Type III (*KCNMA1, SLC4A4*), which we discuss in detail below.

Several of these marker genes, including *ARL15, TSHZ2, MRC1, GRM8*, and *TRPM3*, have not previously been identified as markers for cerebrovascular cell types. We confirmed that these DEGs and corresponding cell types and subtypes were consistent between our *ex vivo* and *post mortem* samples (**Extended Data Fig. 2f**), indicating that they hold across sample types and methodologies. Indeed, the marker genes *VEGFC, GRM8*, and *TRPM3* exhibited protein expression in human cerebrovascular cell types by immunofluorescence (**Fig. 1g**).

### Integrative analysis of human *ex vivo, post mortem*, and mouse cerebrovascular cell types

To evaluate whether the cerebrovasculature expression profiles are conserved across species and platforms, we used canonical correlation analysis^21^ to integrate our 6,873 *ex vivo* and 24,965 *post mortem* human cells with 3,406 cells from C57BL/6 mice^3^. We found that matching cell types co-clustered across species and platforms (**Fig. 2a-b, Extended Data Fig. 3a-b**), indicating broad cell type conservation and enabling us to evaluate differentially expressed genes (DEGs) for each pairwise comparison. We found extensive agreement between the two human datasets, with only 0.7% of differentially expressed genes (*n*=146) between *ex vivo* and *post mortem* samples in each cell type on average (**Supplementary Table 3**), indicating robustness of our findings to platform differences. The very few differentially expressed genes were enriched in mitochondrial genes (101-fold, *p*=6.8e-22), which were found across multiple cell types, possibly reflecting *post mortem* tissue/nuclei degradation. No other functional enrichments were found, indicating that the major functional modularities of human cerebrovascular cell types are preserved despite the *post mortem* interval and subsequent cryopreservation. By contrast, we found extensive differences between species, with 9.3% of genes consistently differentially expressed (*n*=2,002) between human and mouse for each cell type on average (**Extended Data Fig. 3c, Supplementary Table 3**). Human-mouse differentially-expressed genes were highly consistent between *ex vivo* and *post mortem* samples (80% agreement, *p*<2.2e-16). In each cell type, DEGs were strongly enriched for marker genes of that cell type, indicating that cell type identity markers were among those that vary the most between species. In endothelial cells, DEGs were 12-fold enriched for marker genes, in fibroblasts 13.6-fold enriched, in pericytes 15-fold enriched, and in SMCs 8.6-fold enriched.

**Figure 2.**
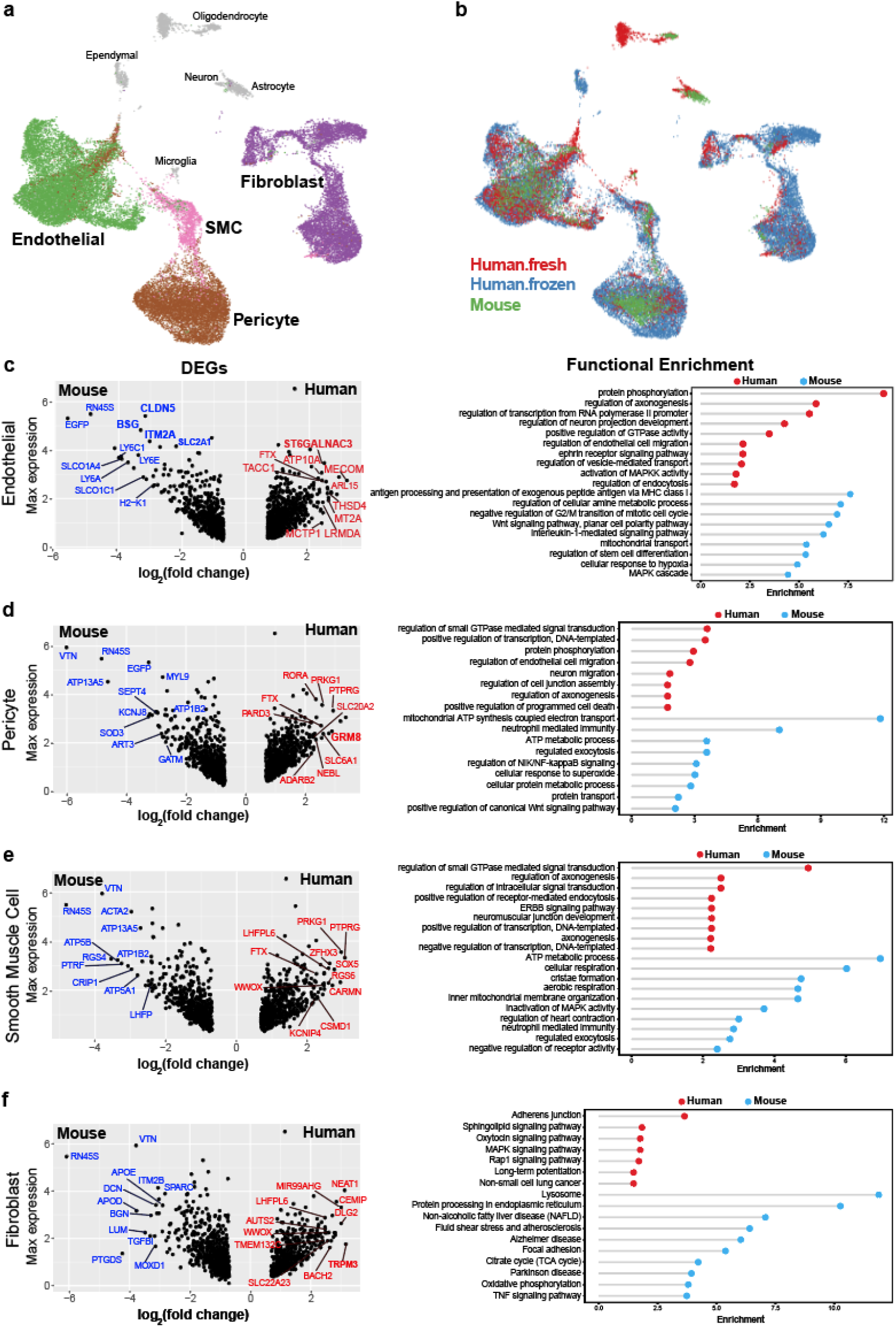
Integrative analysis of human *ex vivo, post mortem*, and mouse cerebrovascular cell types. **a-b**. UMAP visualization of integrated cells from human *ex vivo*, human *post mortem* and mouse, coloring by cell types (**a**) and data source (**b**). All vessel-coupled cells were colored in grey. **c-f**. Differentially expressed genes between human and mouse in endothelial (**c, left**), pericyte (**d, left**), smooth muscle cells (**e, left**), and fibroblast (**f, left**). X-axis represents the log-transformed fold change and y-axis represents the maximal expression level. The top 20 genes are highlighted in blue for mouse and red for human. Genes that were also cell type markers are bolded. **c-f. (right panels)**, the representative functional enriched terms of human-and mouse-specific/highly expressed genes.

Human-mouse DEGs showed cell type-specific gene ontology enrichments that provide important insight into species-specific functional specialization of the cerebrovasculature. Endothelial cells showed 767 human-enriched genes, with broad enrichments in cerebrovascular function, axonogenesis, and endothelial cell migration regulation pathways, and 704 mouse-enriched genes enriched in antigen processing and presentation, metabolic process, and immune response pathways (**Fig. 2c**). Pericytes showed 702 human-enriched genes enriched in signal transduction, neuron migration, and regulation of cell junction assembly pathways, and 781 mouse-enriched genes enriched in regulated exocytosis and ATP metabolic process pathways (**Fig. 2d**). Smooth muscle cells showed 582 human-enriched genes, enriched in GTPase signaling regulation and receptor-mediated endocytosis and other intracellular signaling pathways, and 629 mouse-enriched genes enriched in ATP metabolic process, regulated exocytosis and immune response pathways (**Fig. 2e**). Human-enriched genes included AD-associated^22^ *PICALM*, involved in amyloid-beta clearance through brain vasculature^23^, and muscle differentiation-associated^24^ long-noncoding RNA *CARMN*, suggesting human-specific roles. Fibroblasts showed 478 human-enriched genes, enriched in adhesion junctions, sphingolipid, MAPK, and Rap1 signaling pathways, and 792 mouse-enriched genes, enriched in extracellular matrix organization, endocytosis regulation, and exocytosis regulation pathways (**Fig. 2f**).

### Human-specific endothelium zonation

Endothelial cells show phenotypic zonation^25^ along the arteriovenous axis. Though poorly understood, zonated characteristics arise in diseases, including the observation that small caliber arteries are preferentially affected in cerebral arteriopathies^10^, and that brain capillaries are likely more affected in multiple neurodegenerative disorders^1,6^. To profile high-resolution human endothelial molecular zonation signatures, we developed a continuous quantitative measure of spatial cell positioning using a linear regression model, focusing specifically on our *ex vivo* nuclei. We found 1,802 genes that displayed a pronounced gradient of expression along the arteriovenous axis, including 147 transcription factors and 76 transporters (**Extended Data Fig. 4a,b**), indicating a gradual transcriptional continuum (**Fig. 3a,b**). We found strong conservation between zonation markers of human and mouse arterioles (*VEGFC, BMX, EFNB2*) and capillaries (*MFSD2A, TFRC*), but venule zonation markers differed between species. For example, *SLC38A5* was expressed in both capillaries and venules in human, but only in venules in mouse. *TSHZ2* and *LRRC1* were both venule-zonated in human, but showed no expression in mouse endothelium. We found that several transcription factors previously known to promote general endothelial or arterial fate, including *HEY1* and *GATA2*^*26*^, are also arteriole-zonated, and also reveal here several novel venule-zonated transcription factors, including *TSHZ2, BNC2*, and *ETV6*.

**Figure 3.**
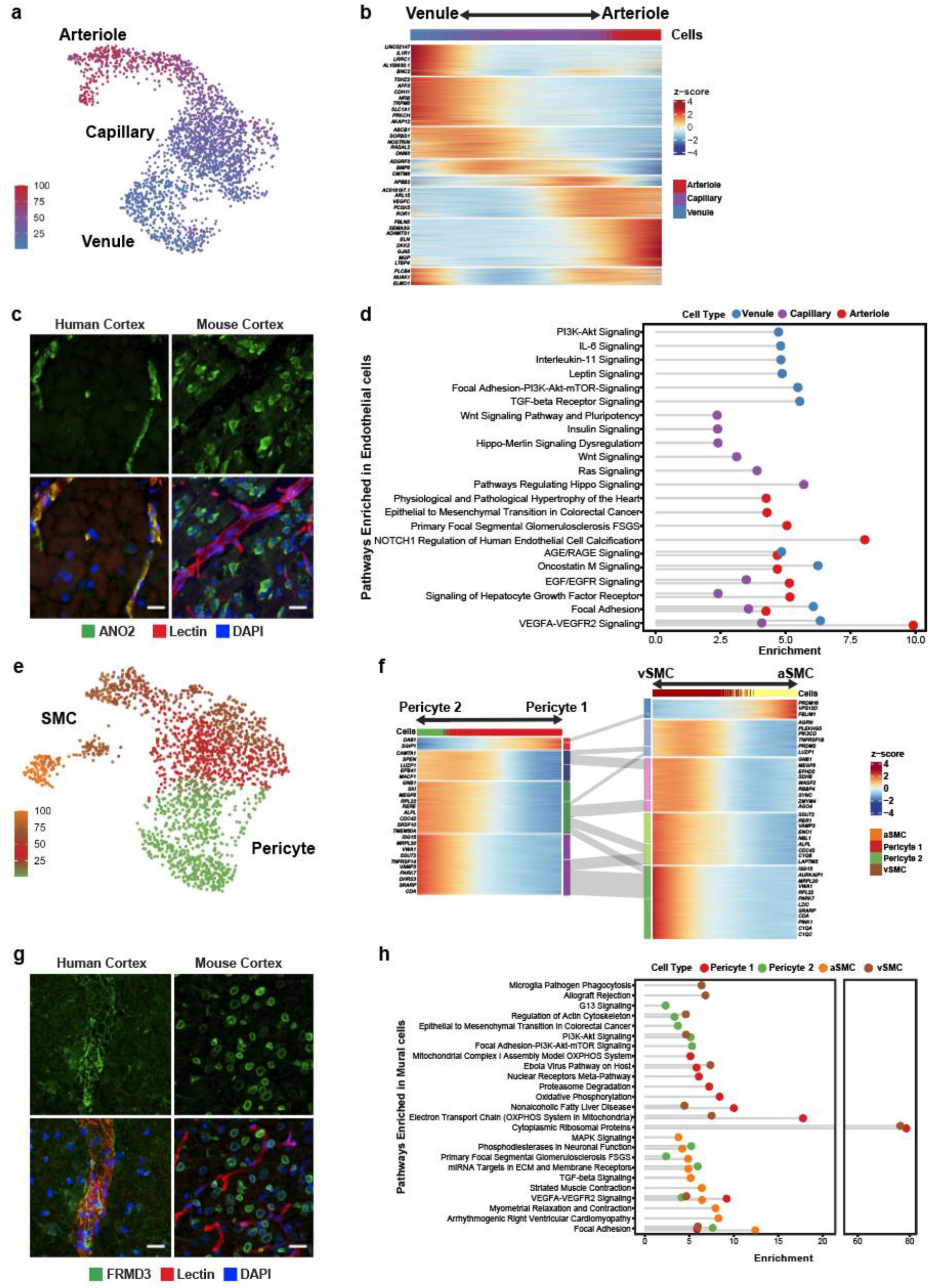
Molecular zonation of human brain endothelial and mural cells. **a**. Zonal gradient of endothelial cell transcriptomes. **b**. Highly expressed genes along the endothelial gradient. **c**. Indirect immunofluorescence of human endothelial marker *ANO2*, and mouse homolog *Ano2*, expression in brain cortex. **d**. Pathway analysis along endothelial zones. **e**. Zonal gradient of mural cell transcriptomes. **f**. Highly expressed genes along the mural gradients. **g**. Indirect immunofluorescence of the human mural marker *FRMD3*, and mouse homolog *Frmd3*, expression in brain cortex. **h**. Pathway analysis along mural zones. Scale bar, 20μm.

We experimentally validated the zonated expression of several genes using indirect immunofluorescence staining of freshly resected cortical tissue. We confirmed human-specific capillary/venule zonation for *ANO2* (**Fig. 3c**), which lacks endothelial cell expression in mouse brain^13^, and human-specific venule zonation for *TSHZ2* (**Extended Data Fig. 5a**), and human-mouse conserved^3^ arteriole zonation for *VEGFC*. We also found human-specific expression for zinc-binding metallothioneins *MT1E* and *MT2A* (**Extended Data Fig. 5b,c**) that show astrocyte expression but no vascular expression in mouse.

Differentially-zonated genes were enriched in distinct functional gene ontologies and pathways (**Fig. 3d, Extended Data Fig. 5d**). For example, venule-zonated genes were enriched in IL-6 and IL-11 signaling pathways, consistent with the central role of venules in leukocyte adhesion and cytokine release^27,28^. Arterial-zonated genes were enriched in Notch signaling, consistent with the role of Notch mutations in small-vessel arteriopathies^10^. These results highlight that while zonated functional organization is a conserved characteristic across species, the human cerebrovasculature exhibits a species-specific gene expression pattern.

### Human brain mural cells exhibit discrete molecular zonation

Unlike endothelial cells, mural cells exhibited two distinct and separate transcriptional gradients for pericytes and SMCs, independent of their position along the arteriovenous axis (**Fig. 3e**). Though heterogeneity in mural cells is known, these discrete gradients of gene expression were not previously observed in mice^3^. Altogether, we identified 1,820 zonated genes in pericytes (**Extended Data Fig. 6a,b**) and 2,756 zonated genes in SMCs (**Extended Data Fig. 7a,b**) which display distinguishable expression patterns (**Fig. 3f**). Interestingly, we found that the zonated genes in SMCs and pericytes are significantly shared based on their expression patterns. For example, the genes highly expressed in Pericyte 2 were enriched in vSMCs. In contrast, genes highly expressed in Pericyte 1 were highly expressed in aSMCs (**Fig. 3f, Extended Data Fig. 7c**).

We found zonated expression of several apolipoproteins in vSMCs, including *APOD, APOE*, and *APOO*. Interestingly, failure of amyloid-beta clearance in the perivenous space is thought to contribute to CAA and deficits of perivenous drainage in Alzheimer’s disease^29,30^. Given the localized recruitment of immune cells to venules, the zonated expression of apolipoproteins, in particular *APOE*, by vSMCs suggests a zonated functional role of amyloid-beta clearance at the level of venules. In addition, our analysis revealed the expression of many novel genes in human mural cells. In pericytes, we validated the zonated expression of metabotropic glutamate receptor 8, *GRM8* (**Extended Data Fig. 8a**). For SMCs, we validated the zonated expression of *MRC1, FRMD3, SLC20A2*, and *SLC30A10* (**Fig. 3g, Extended Data Fig. 8b,d-e**). Importantly, these SMC-specific genes exhibited expression predominantly wrapped around large caliber microvessels, but not in perfect co-localization with the endothelial marker lectin. Furthermore, all genes exhibited expression patterns in non-vascular cell types within the mouse posterior cortex, demonstrating the species-specific expression in human mural cells.

Lastly, our gene ontology and pathway analyses indicated that highly expressed genes in Pericyte 2 and aSMCs were significantly enriched in vascular smooth muscle contraction, cardiac muscle cell action potential, and cell-cell adhesion terms. Likewise, Pericyte1 and vSMC highly expressed genes were significantly enriched in VEGF signaling pathway, focal adhesion, ABC transporters, immune responses (antigen processing and presentation and phagocytosis) terms (**Fig. 3h, Extended Data Fig. 8c**). The pathways enriched in Pericyte 1 more closely resembled vSMCs, whereas Pericyte 2 more closely resembled aSMCs, suggesting functional similarities between aSMC/Pericyte 2 and vSMC/Pericyte 1, and likely reflecting a diversity in human pericyte function depending on their proximity to arterioles or venules.

### Three types of perivascular fibroblasts in human cortex

Endothelial cells, mural cells, and astrocytes are notable cellular components of the brain vasculature; however, recent studies in mice and zebrafish have demonstrated a class of perivascular fibroblasts as being integral for vascular structure^3,13,31^. In particular, these perivascular fibroblasts, previously referred to as stromal cells, are known to express collagens and laminins, which are essential components of the extracellular matrix^32^ and contribute to fibrotic scar formation after brain or spinal cord injury^33–36^. Distinct fibroblast subtypes have been determined to be present in the mouse cerebrovasculature^3,37^; however, the limited number of fibroblasts have prevented thorough characterization of their distinct transcriptional profiles and function.

In our sub-clustering analysis of vascular cell types, we uncovered three distinct subtypes of perivascular fibroblasts (**Fig. 4a**), two of which were consistent with those previously identified in mouse (Types I and II). Type III fibroblasts shared expression of the mouse arachnoid barrier cell marker^37^, *SLC47A1*; however, several other markers of arachnoid barrier cells are not expressed in our human Type III fibroblasts. As we did not expect to have arachnoid barrier cells present in our precise human *ex vivo* tissue resections, the human Type III fibroblasts revealed by our analysis likely constitute a novel human fibroblast subtype not previously described. Nevertheless, all subtypes expressed highly specific sets of genes (**Fig. 4b**), some of which are also expressed in mouse brain fibroblasts, though subtype-specific expression had not been previously assessed. Here, we confirm by indirect immunofluorescence staining the expression of *FBLN1, CEMIP*, and *KCNMA1*, as markers for Type I, II, and III fibroblasts, respectively (**Fig. 4c**).

**Figure 4.**
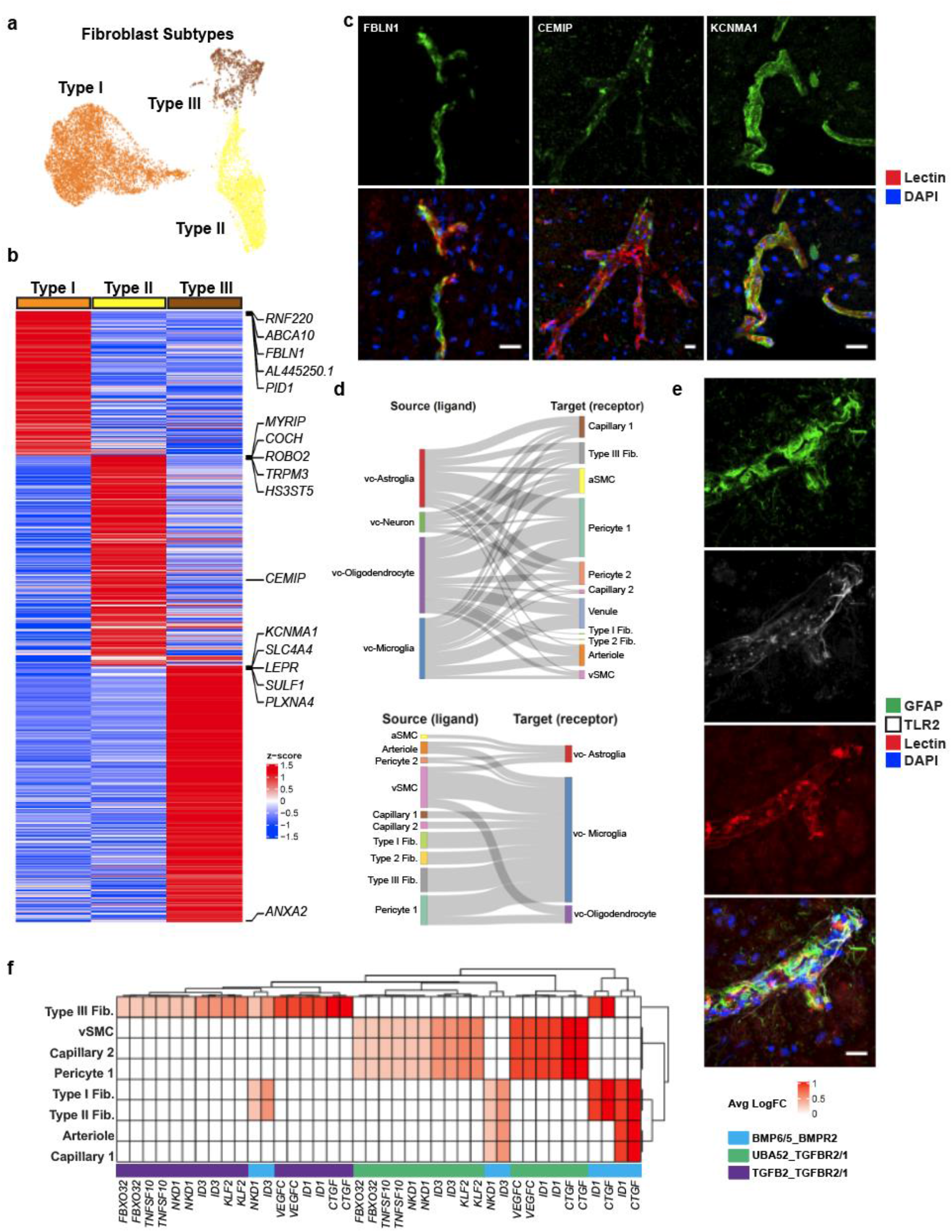
Cell types coupled to the human cerebrovasculature. **a**. UMAP of integrated perivascular fibroblast subtypes from human *ex vivo* and *in silico* sorted brains. **b**. Highly expressed genes in fibroblast subtypes. **c**. Indirect immunofluorescence of fibroblast markers *CEMIP, TRPM3*, and *KCNMA1* expression. **d**. Cell-cell interaction Sankey plots between vascular cell types (endothelial, mural, fibroblast) and canonical cell types (astrocyte, neurons, microglia, oligodendrocytes). **e**. Indirect immunofluorescence of *GFAP/TLR2* expressing astrocytes coupled with the cerebrovasculature. **f**. Heatmap of ligand-receptor pairings in vascular cells (ligand) and microglia (receptor) and downstream target. Scale bar, 20μm.

Pathway and gene ontology enrichment analyses revealed distinct functional roles for each subtype (**Extended Data Fig. 9a,b**). While all subtypes expressed genes involved in VEGF-VEGFR2 signaling, suggesting a close interaction with the brain vasculature, based on their gene expression signatures, Type I fibroblasts appear to be the main subtype involved in ECM organization. Type II and Type III showed greater significance in pathways related to cell fate, with Type III showing expression of various growth factors, including *VEGFA*. Pseudotime analysis of *ex vivo* fibroblasts revealed two gradients of gene expression from Type I to Type II and to Type III fibroblasts, separately (**Extended Data Fig. 9c**). Interestingly, the Type I to Type II trajectory was continuous with a subpopulation of pericytes, Pericyte 2 (**Extended Data Fig. 9d**), suggesting a lineage from Type I to Type II to pericytes and consistent with a recent study demonstrating the stem cell potential of fibroblasts to differentiate into pericytes^31^. Therefore, Type II fibroblasts likely represent an intermediate state exhibiting a greater mural cell transcriptional phenotype.

### Vasculature-coupled (VC-) classical brain cell types

Neurons and glia are known to interact extensively with the vasculature. Endothelial cells have been shown to synthesize BDNF as a guidance cue for neuronal precursors to migrate along the vasculature in the adult forebrain^38^. Similarly, astrocytes have been shown to provide complementary scaffolds for neuroblasts^39^, and microglia migrate postnatally along the vasculature to occupy perivascular regions devoid of astrocytic end-feet^40^. How these particular subsets of neuronal and glial cell types are coupled to the vasculature and how their transcriptional profiles differ from non-coupled cells remain unanswered questions.

In our *ex vivo* samples, as expected by physical BVE protocol co-purification, we found vasculature-coupled (vc) neuronal and glial cells with distinct expression profiles from the corresponding canonical cell types (**Fig. 1c**). Despite showing features of both canonical and vascular cells, we confirmed vasculature-coupled cells do not stem from ambient RNA or doublets, as they were unaffected by two rounds of doublet exclusion at the cluster and cell levels and ambient RNA removal steps (**Extended Data Fig. 10a-b**). Hundreds of specifically expressed genes in these vasculature-coupled cells, when compared to canonical cell types, included significant enrichment in the regulation of angiogenesis, metal ion transport, and protein localization terms (**Extended Data Fig. 10c, Supplementary Table 4**). In particular, vc-astrocytes and vc-microglia showed increased expression of genes involved in the negative regulation of cell migration and cell junction assembly, suggesting these cell types exhibit more stationary behavior (e.g. less dynamic processes) when coupled to the vasculature (**Extended Data Fig. 10d**). We validate that a small number of *NeuN/RBFOX3*-expressing neurons can be found immediately adjacent, seemingly coupled, to blood vessels (**Extended Data Fig. 10e**).

To reveal mechanisms of communication between vasculature-coupled and vascular cell types, we computationally integrated cell type-specific ligand, receptor, and downstream target gene expression in a statistical permutation framework (see Methods). We found that vc-astrocytes, vc-oligodendrocytes, and vc-microglia all interact extensively with Pericyte 1 cells, as a major signal-receiving vascular cell type (**Fig. 4d**). We confirm *TLR2* enrichment in vc-astrocytes by indirect immunofluorescence staining to demonstrate that a subset of *GFAP*-expressing astrocytes in proximity to the vasculature express *TLR2*, whereas *GFAP*-expressing astrocytes not in proximity to blood vessels do not express *TLR2* (**Fig. 4e, Extended Data Fig. 10f**).

Lastly, we found vc-microglia receive signals from many vascular cell types, including vSMCs, all fibroblast subtypes, and pericytes, indicating that they may act as central hubs for vasculature interactions. Predicted receptors *TGFBR1, TGFBR2*, and *BMPR2* were highly expressed in vc-microglia, indicating they may act as mediating receptors. These receptors activate downstream targets *ID1, VEGFC*, and *ID3*, which were also highly expressed in vc-microglia (**Fig. 4f**).

### Huntington’s disease cerebrovascular dysfunction signatures

Recent studies have noted cerebrovascular abnormalities across many neurodegenerative diseases, with BBB integrity breakdown often preceding more disease-specific pathological features^7,41–43^. In particular, Huntington’s disease (HD), an incurable monogenic neurodegenerative disorder caused by CAG trinucleotide repeat expansion in the huntingtin (*HTT*) gene^44^, has been associated with various abnormalities to the cerebrovasculature and BBB in both human patients and mouse models, including increased BBB permeability, increased small vessel density, altered morphology of blood vessels, and altered cerebral blood volume^42,45–52^, but the molecular bases of these alterations are currently not well understood.

As proof-of-principle of the utility of our annotations to study cerebrovascular cells in the context of disease, we used our high-resolution cerebrovasculature cell type annotations to investigate vascular cell gene expression alterations in HD, to annotate 2,198 cells from our previously-published *post-mortem* neostriatal HD samples^18^, and study their gene expressions hanges in HD. Illustrating the utility of the reference data produced here for properly annotating human BBB cell types, we corrected the annotation 895 fibroblasts that we had previously^18^ incorrectly labeled as mural cells (**Fig. 5a, Extended Data Fig. 11a-b**). We also profile here an additional 1,747 cells from HD grade 1-4 and control cells purified with the BVE protocol. Highlighting the importance of our vasculature-coupled cell signatures, we found that one previously-uncharacterized subcluster of astrocytes and two previously-uncharacterized subclusters of microglia showed our vasculature-coupled signatures, enabling us to annotate 2,758 vc-astrocytes and 1,004 vc-microglia in our HD samples (**Extended Data Fig. 11c-e**). The number of vc-astrocytes and vc-microglia were significantly increased in HD (**Fig. 5b**), suggesting that astrocytes and microglial cells associate more closely with the vasculature in HD.

**Figure 5.**
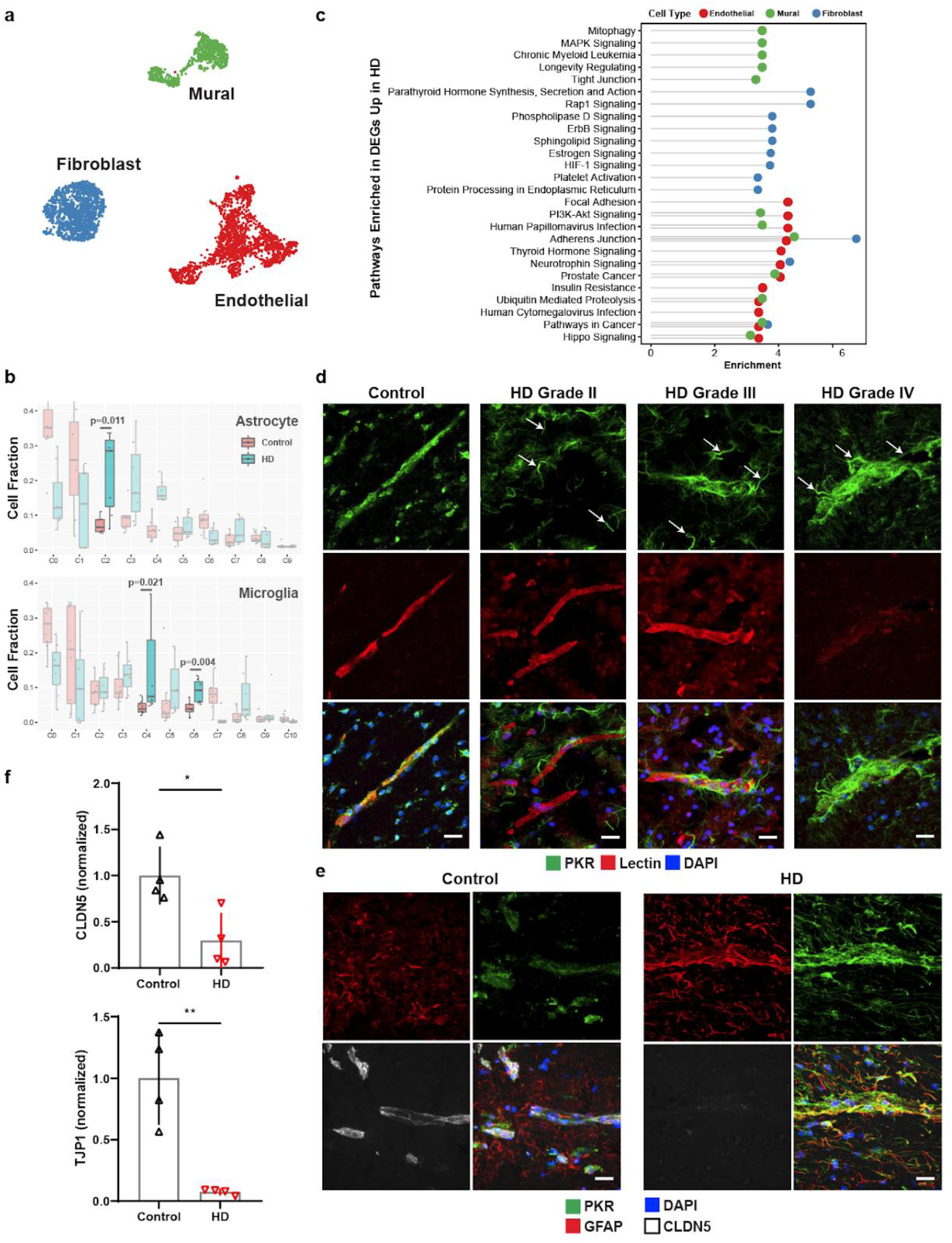
Innate immune activation related to cerebrovascular dysfunction in Huntington’s disease (HD). **a**. UMAP of integrated cerebrovasculature cells in *post mortem* control and HD human patients. **b**. Cell fraction analysis of astrocyte and microglia subclusters; statistics shown for highlighted clusters using the Wilcoxon rank-sum test. **c**. Pathway analysis of the top 10 enriched upregulated pathways in HD endothelial, mural, and fibroblasts cells. **d**. PKR immunostaining in perivascular glial processes across various HD grades. PKR is normally detected in the vasculature in control samples; arrows indicate PKR-immunopositive vasculature-coupled glial processes that become apparent in HD samples only. **e**. PKR immunostaining co-localizes with GFAP and engulfs blood vessels with low CLDN5 expression. **f**. Western blot quantification for tight junction proteins CLDN5 and TJP1. Scale bar, 20μm. WB: two-tailed t test, \**p* < 0.01, \*\**p* < 0.001.

We found 4,698 differentially expressed genes (DEGs) between HD and controls for endothelial, mural, and fibroblast cell types (**Supplementary Table 5**), some of which are known to be dysregulated in neurodegenerative conditions, including *ABCB1, ABCG2, SLC2A1* downregulation in endothelial cells, and *PDGFRB* downregulation in mural cells. We also found significant endothelial down-regulation of *MFSD2A*, a lipid transporter that is expressed in brain endothelial cells and is important in restricting caveolae-mediated transcytosis at the BBB^53–55^, suggesting its downregulation may underlie increased vesicular trafficking and BBB leakage in HD, as is observed in aged mice^56^. In addition, endothelial HD cells showed upregulation of sprouting angiogenesis, endothelial cell migration, and vascular endothelial growth factor signaling genes (**Supplementary Table 5**), changes which could be responsible for the increased vessel density that is noted in HD^42^. We also predicted upstream regulators of differentially expressed genes using chromatin enrichment analysis (ChEA, Methods). We found that *TCF4*, a master regulator of the *Wnt* pathway, was a top predicted regulator of upregulated genes in HD endothelial cells, and was itself upregulated in endothelial cells (**Supplementary Table 5, Extended Data Fig. 11f**). Consistent with this finding, the Wnt signaling pathway is upregulated in brain microvasculature endothelial cell (iBMEC) studies utilizing HD patient-derived iPSC cells^8^.

HD-upregulated genes in endothelial cells were enriched for many viral-responding innate immune activation genes, including the Human Papillomavirus and Cytomegalovirus Infection terms (**Fig. 5c**), complementing our previously observed innate immune activation in human HD neostriatal spiny projection neurons^18^, and which is of interest given previously-observed links between BBB dysregulation and innate immune activation^57–59^. Indeed, several key initiators and mediators of innate immune activation were upregulated in endothelial cells, including *HSPH1, IL1RAPL1, IL3RA, IL4R, IL6R, IKBKB, IRF2/3, IFNAR1, NFKBI*, and *STAT3* (**Supplementary Table 5**). We found that innate immune activation genes were also significantly upregulated in HD for both vc-astrocytes and vc-microglia, including *HSPH1, IL6R, IRF3, NFKBI, NFKBIA*, and *TRAF3* (**Supplementary Table 6**). To validate this upregulation of innate immune signaling in vc-astrocytes and vc-microglia, we performed indirect immunofluorescence staining for the innate immune sensor PKR in HD and control samples from both human and mouse model tissue. In control tissue, some level of PKR expression is normally detected in the cerebrovascualture. We observed a strong PKR protein upregulation in what appeared to be glial processes that engulfed blood vessels, in both human HD (**Fig. 5d**) and R6/2 HD mouse model samples (**Extended Data Fig. 11g**). These engulfed blood vessels were devoid, or exhibited low levels of, the BBB tight junction protein CLDN5, and the engulfing processes were immunopositive for astrocyte marker GFAP, suggesting a correlation between endothelial and glial cell innate immune activation and reduction of endothelial tight junction protein levels (**Fig. 5e, Extended Data Fig. 11g**). We further confirmed a significant reduction to overall levels of the BBB tight junction proteins CLDN5 and TJP1 by Western blotting (**Fig. 5f, Extended Data Fig. 11h**). Since downregulation of CLDN5 and TJP1 levels are well known to lead to the loss of BBB integrity, taken together these data provide evidence for a co-occurance of activation of innate immune signaling in endothelial cells/vasculature-coupled glial cells and the loss of BBB integrity that has been noted in HD and HD mouse models.

## Discussion

Interrogating cell types of the cerebrovasculature in humans at a molecular level has been challenging due to a lack of a high-throughput methodology to capture these cell types for genome-wide gene expression analyses. Species-specific patterns of gene expression between mouse and human brain microvessels have been noted, but the lack of cellular specificity and resolution have prevented a thorough mapping of their molecular composition^60^. Here, by developing independent experimental and *in silico* sorting methodologies for the enrichment of brain microvessels that are compatible with snRNA-seq, we catalog the transcriptional profiles of thousands of single nuclei comprising the human cerebrovasculature. Included in our profiles are those generated from fresh human cortex surgical resections that eliminate any possibility of transcriptional alterations associated with the *post mortem* interval. These profiles reveal previously unknown human specific characteristics of the cerebrovasculature. We observe that vascular zonation is overall an evolutionarily conserved phenomenon when comparing mice to humans. However, although arteriole-zonated genes were found to be consistent across species, venule-zonated genes in humans were divergent from those in mouse. Furthermore, across species we observed similar degrees of diversity and specificity in the gene expression patterns of mural cells, which until now have remained ill-defined. We validate the expression of novel marker genes for both pericytes and smooth muscle cells, confirming the cell type-specificity of expression at the protein level. In addition, we identify three subtypes of perivascular fibroblasts and a subset of classical cell types that are closely coupled to the cerebrovasculature. By analyzing their gene expression profiles and cell-cell interactions, we determined a closely tied network of receptor-ligand interactions which have potential implications on their functional roles with respect to the cerebrovasculature.

Understanding the normal cellular and molecular characteristics of the cerebrovasculature is crucial for studying mechanisms of dysfunction (e.g., BBB breakdown and neurodegeneration). In our study, we also conducted cell type-specific differential gene analysis of cerebrovascular cells in the context of HD to elucidate potential mechanisms of BBB dysfunction. In addition to the previously studied Wnt signaling pathway^8^, our work reveals the activation of the innate immune signaling, which we have recently reported to occur in striatal spiny projection neurons^8,18^, as also occurring in endothelial cells and vasculature-coupled astroglial and microglial cells in the HD brain. Our work also reveals that this innate immune activation co-occurs with loss of BBB tight junction protein expression, suggesting that blockade of innate immune activation in the HD brain could prevent loss of BBB integrity.

Our study demonstrates the sophisticated nature of cell types comprising the human cerebrovasculature. These human-specific profiles will be an invaluable source for both understanding basic characteristics of the cerebrovasculature and mechanisms of dysfunction in pathological states.

## Methods

### Animal Usage

All animal experiments were approved by the MIT Committee on Animal Care. Mice were grouped housed with food and water provided *ab libitum* on a standard 12h light/12h dark cycle. 6-week old male C57BL/6J wild-type mice (Jackson Laboratories stock #000664) were used for snRNA-seq and immunofluorescence experiments. 9-week old male B6CBA-Tg(HDexon1)62Gpb/1J mice (CAG repeat length 160 ± 5; Jackson Laboratories stock # 002810) and non-carrier controls were used for R6/2 experiments. No prior procedures were performed on any animal prior to experiments. All brain dissections were performed on dry ice after cooling the head in liquid nitrogen for 3 sec. Dissected whole brain or posterior cortex were subsequently flash frozen in liquid nitrogen and stored at −80°C until further use.

### Human Tissue

All resected *ex vivo* human tissue was obtained fresh or fresh-frozen from Boston Children’s Hospital through the Repository Core for Neurological Disorders. Upon surgical resection, tissue was examined by a licensed neuropathologist and allocated for clinical or research purposes. Non-pathologically deemed tissue was subsequently stored at −80°C until further use. All selected patient samples were between 11-22 years of age and had a primary diagnosis of medically refractory epilepsy with no known genetic mutations (i.e., spontaneous epilepsy). Cases with known arteriovenous malformations were also excluded from the study. 8 human *post mortem* tissue from HD and 8 age-matched unaffected controls were acquired through the NIH NeuroBioBank or the University of Alabama at Birmingham. 12 unaffected controls without VCI or dementia were obtained from the Religious Orders Study and Rush Memory and Aging Project (ROSMAP, approved by an Institutional Review Board (IRB) of Rush University Medical Center, ROSMAP resources can be requested at https://www.radc.rush.edu). These included 7 brain regions: prefrontal cortex, mid-temporal cortex, angular gyrus, entorhinal cortex, thalamus, hippocampus and mammillary body.

### Indirect Immunofluorescence

Brain tissue was harvested from the HD R6/2 model and the control mice following transcardial perfusion with 4% PFA in 1X PBS. Human and mouse brain tissue samples were cryoprotected and cryosectioned onto glass slides at 20 µm thickness. Sections were fixed for 10 min using cold acetone (human only) under gentle agitation, washed with 1X TBS, permeabilized with 1X TBS-T (1X TBS with 0.05% Tween20), and blocked with blocking buffer (2% heat-inactivated donkey serum, 0.1% fish gelatin in 1X TBS-T) for 1 hr at room temperature. Slides were subsequently incubated with primary antibody in blocking buffer at specified dilutions (**Supplementary Table 7**) overnight at 4°C. Slides were then washed with 1X TBS-T, incubated with secondary antibody (1:5,000 dilution of specified fluorophore as described in **Supplementary Table 7**) in blocking buffer for 1 hr at room temperature, washed again with 1X TBS-T followed by 1X TBS, stained with DAPI, and mounted with ProLong Gold antifade mounting media (ThermoFisher Scientific, Waltham MA). For *post mortem* human tissue sections, auto-fluorescent signal arising from endogenous lipofuscin was quenched with a 30-second exposure to TrueBlack® Lipofuscin Autofluorescence Quencher (Biotium, Fremont, CA) and washed with 1X-TBS prior to mounting. A Zeiss LSM700 confocal microscope (Carl Zeiss AG, Oberkochen, Germany) with a 20X and 40X objective lens was used for imaging. All image processing was done using ImageJ software.

### Blood Vessel Enrichment for Single Nuclear RNA Sequencing

BVE protocol was adapted from Lee et al.^14^ and Mathys et al.^15^. All procedures were performed on ice. *Ex vivo* human, *post mortem* human, and dissected mouse cortical tissue were homogenized in 5mL of MCDB 131 media containing 0.5% (wt/vol) endotoxin-, fatty-acid- and protease-free BSA and 10 U ul^−1^ units of recombinant RNase Inhibitors using 10 strokes with the loose pestle followed by 10 strokes with the tight pestle in a 7mL KIMBLE Dounce tissue grinder. Homogenized tissue was transferred into 15mL conical tubes and centrifuged at 2,000 x g for 5 min at 4°C using a fixed angle rotor. Cell pellet was resuspended in 2mL of 15% (wt/vol) 70-kDa dextran solution (in 1X PBS + 10 U ul^−1^ recombinant RNase Inhibitors) and ultracentrifuged at 10,000 x *g* for 15 min at 4°C using a swing-bucket rotor. Resultant pellet or fresh frozen tissue was homogenized in 700µL of Homogenization Buffer (320 mM sucrose, 5 mM CaCl_2_, 3 mM Mg(CH_3_COO)_2_, 10 mM Tris HCl pH 7.8, 0.1 mM EDTA pH 8.0, 0.1% NP-40, 1 mM β-mercaptoethanol, and 0.4 U µl^−1^ SUPERase In RNase Inhibitor (ThermoFisher Scientific, Waltham MA)) with a 2mL KIMBLE Dounce tissue grinder (MilliporeSigma, Burlington MA) using 10 strokes with the loose pestle followed by 10 strokes with the tight pestle. Homogenized tissue was filtered through a 40µm cell strainer and mixed with 450µl of working solution (50% OptiPrep density gradient medium (MilliporeSigma, Burlington MA), 5 mM CaCl_2_, 3 mM Mg(CH_3_COO)_2_, 10 mM Tris HCl pH 7.8, 0.1 mM EDTA pH 8.0, and 1 mM β-mercaptoethanol). The mixture was then slowly pipetted onto the top of an OptiPrep density gradient containing 750µL of 30% OptiPrep Solution (134 mM sucrose, 5 mM CaCl_2_, 3 mM Mg(CH_3_COO)_2_, 10 mM Tris HCl pH 7.8, 0.1 mM EDTA pH 8.0, 1 mM β-mercaptoethanol, 0.04% NP-40, and 0.17 U µL^−1^ SUPERase In RNase Inhibitor) on top of 300µL of 40% OptiPrep Solution (96 mM sucrose, 5 mM CaCl_2_, 3 mM Mg(CH_3_COO)_2_, 10 mM Tris HCl pH 7.8, 0.1 mM EDTA pH 8.0, 1 mM β-mercaptoethanol, 0.03% NP-40, and 0.12 U µl^−1^ SUPERase In RNase Inhibitor) inside a Sorenson Dolphin microcentrifuge tube (MilliporeSigma, Burlington MA). Nuclei were pelleted at the interface of the OptiPrep density gradient by centrifugation at 10,000 x *g* for 5 min at 4°C using a fixed angle rotor (FA-45-24-11-Kit). The nuclear pellet was collected by aspirating ~100µL from the interface and transferring to a 2.5mL low binding Eppendorf tube. The pellet was washed with 2% BSA (in 1X PBS) containing 10µL^−1^ SUPERase In RNase Inhibitor. The nuclei were pelleted by centrifugation at 300 x g for 3 min at 4°C using a swing-bucket rotor (S-24-11-AT). Nuclei were washed two times with 2% BSA and centrifuged under the same conditions. The nuclear pellet was resuspended in ~100µL of 2% BSA.

### Quantitative Real-Time PCR

Mouse posterior cortex was disrupted for RNA isolation using the TissueLyser (QIAGEN, Hilden, Germany) for 2 x 2 min at 20 Hz as recommended by the manufacturer. Enriched blood vessel pellets from the BVE protocol were disrupted using QIAGEN Buffer RLT. RNA was then isolated using the RNeasy Lipid Tissue Mini Kit (QIAGEN, Hilden, Germany). For qRT-PCR, the TaqMan Universal Master Mix (ThermoScientific, Rockford, IL) was used, and PCR reactions were run on a StepOnePlus system (ThermoScientific, Rockford, IL).

### Western Blotting

Human *post mortem* caudate nucleus tissue samples from HD and age-matched unaffected controls were homogenized in 1mL of RIPA lysis buffer (ThermoScientific, Rockford, IL) containing a protease inhibitor cocktail (MilliporeSigma, Burlington MA). ~10μg of protein in 1X LDS were loaded onto 4-12% Bis-Tris gels and run using MOPS Running Buffer at 175V for 1 hour. Protein samples were then transferred onto PVDF membranes (Bio-Rad, Hercules, CA) using 10% methanol in 1X transfer buffer at 50V for 1.5 hours.

Membranes were then washed in 1X TBS-T (1X TBS with 0.05% Tween20), blocked with 5% milk for 1 hour at room temperature, and subsequently incubated with primary antibodies (**Supplementary Table 7**) in 5% milk at 4°C overnight under gentle agitation. Membranes were then washed again with 1X TBS-T, incubated with HRP secondary antibodies for 1 hour at room temperature, and applied with substrate (Pierce ECL Plus) for chemiluminescent detection. Immunoblots were quantified using ImageJ software. Two-tailed t-test was used for statistical analysis.

### Single Nuclear RNA Sequencing and Analysis

Droplet-based snRNA sequencing libraries were prepared using the Chromium Single Cell 3′ Reagent Kit v3 (10x Genomics, Pleasanton CA) according to the manufacturer’s protocol and sequenced on an Illumina NovaSeq6000 at the MIT BioMicro Center. Raw sequencing reads were aligned to the pre-mRNA annotated Mus musculus reference genome version GRCm38 or Homo sapiens reference genome version GRCh38 and counts were estimated using Cellranger 3.0.1 (10x Genomics, Pleasanton CA). The generated cell-by-gene unique molecular identifier (UMI) count matrix was analyzed using the Seurat R package v.3.2.0^21^. We only kept the cells expressing at least 500 genes and genes with expression in at least 50 cells. The cells were also filtered by the maximum of 8000 expressed genes and of 10% mitochondrial genes. The UMI counts were then normalized for each cell by the total expression, multiplied by 10000, and log-transformed. We used Seurat’s default method to identify highly variable genes and scale data for regressing out variation from UMI and mitochondrial genes. The scaled data with variable genes were used to perform principal component analysis (PCA). The top 30 principal components were chosen for further analysis, including clustering to identify cell populations. UMAPs were calculated in the Seurat R package using the top 30 PCs and min_dist=0.75. Harmony was used to perform batch-effect correction.

### Doublet Removal

To remove the potential doublets in the dataset, we first used DoubleFinder with the parameter of 7.5% doublet formation rate based on the recommendation of 10x Genomics at the single-cell level to identify the most likely doublets^61^. Then, for each cluster, the cluster that highly expressed genes could be significantly enriched in two or more markersets of cell types was also considered as the doublet cluster and removed from further analysis.

### *In Silico* Sorting Approach

We first annotated the cell type for each cluster based on the canonical markers of vascular cell types and then calculated cell type score for each cell based on the average expression of a set of vascular markers^62^. The cells with the specific high score of vascular cell types were kept for further integrative analysis.

### Cell Type Annotation and Marker Genes Identification

To annotate the cell type for each cluster, we identified the marker genes using the Wilcoxon rank-sum test by comparing one cluster with the left. Next, we checked the canonical markers (excitatory neuron: *NRGN, SLC17A7*, and *CAMK2A*; inhibitory neuron: *GAD1* and *GAD2*; astrocyte: *AQP4* and *GFAP*; oligodendrocyte: *MBP, MOBP*, and *PLP1*; microglia: *CSF1R, CD74*, and *C3*; OPC: *VCAN, PDGFRA*, and *CSPG4*; endothelial: *FLT1* and *CLDN5*; pericyte: *AMBP*) in each cluster to determine the cell type. Finally, we performed gene set enrichment analysis by testing the significance of overlapping genes between marker genes that we identified and the published marker gene sets^62^ to further confirm the cell type of each cluster.

### Integrative Analysis of Human Fresh, Frozen and Mouse Single Cell RNAseq Datasets

The homolog genes of human and mouse were kept for integration. Canonical correlation analysis in Seurat^21^ was used to integrate human snRNA-seq data from fresh and frozen samples and mouse scRNA-seq data^3^. To compare the difference between human fresh and frozen samples, we applied MAST in R to identify the differentially expressed genes by considering age and sex as the covariates^63^. We used the Wilcoxon rank-sum test to identify the differentially expressed genes between human and mouse data (two comparisons: human fresh vs mouse, human frozen vs. mouse) and tested the significance of agreement between two comparisons by Fisher’s exact test.

### Functional Enrichment Analysis

Enrichr in R^64–66^ was used to perform functional enrichment analysis based on the following databases: Gene Ontology 2018^67,68^, KEGG/WikiPathways 2019 Human, and ChIP Enrichment Analysis 2016 (ChEA). FDR < 0.05 was used as a threshold to select the significant enrichment.

### Endothelial and Mural Zonation Analysis

The cell orders along the endothelial and mural zonation were determined by the pseudotime analysis using Monocel3^69^. We next built a quadratic linear regression model to identify the zonation-related genes, smoothed the gene expression along the predicted zonation axis by fitting a smoothing spline in R, and clustered those genes into eight distinct expression patterns.

### Cell-Cell Interaction Analysis

We build a computational framework to predict the interaction between vascular cell types and vasculature-coupled cell types. Specifically, we first integrated the information of the ligand-receptor pairs from Ramilowski et al.^70^ and the target genes from KEGG 2019 signaling pathways. We then filtered the ligand, receptor and target genes by the cell type-specific expression. For each pair of cell types, we calculated a score for each ligand-receptor pair to evaluate the probability of interaction between two cell types and mediated path by the ratio of number multiplication of cells with the expression of ligand and receptor to the universal set. To test the significance of each ligand-receptor pair, we permuted the cell ID in the gene expression matrix and calculated the *p*-value of the score.

### Differential Gene Expression Analysis

We identify the differentially expressed genes between control and HD samples using a so-called multiresolution method in ACTIONet^71^. Briefly, the pseudobulk gene expression matrix for each sample is generated based on a number of multiresolution bins (by default is 25). The single-cell variance in pseudobulk data is also considered as a covariate when Limma is applied for differential analysis. In addition, age, sex, and PMI are also controled as covariates in the model. Genes with FDR < 0.05 and Log_2_FoldChange > 0.05 were used for subsequent functional enrichment analysis as described above.

## ACKNOWLEDGMENTS

This research was supported in part by the Intellectual and Developmental Disability Research Center (funded by NIH U54 HD090255) and Rosamund Stone Zander Translational Neuroscience Center at Boston Children’s Hospital, a Picower Institute Innovation Fund Awards and a Walter B. Brewer (1940) MIT Fund Award (to M.H.), and NIH AG054012, AG058002, AG062377, NS110453, NS115064, AG067151, AG062335, MH109978, MH119509, HG008155 and Cure Alzheimer’s Fund CureAlz-CIRCUITS (to M.K.). ROSMAP was supported by NIA grants P30AG10161, R01AG15819, R01AG17917, and U01AG61356. We thank Audrey Helen Effenberger with assistance in graphics design; S. Sebastian Pineda for assistance with differential gene expression analysis; Preston Ge for assistance with the Western blot experiments; and Li-Lun Ho, Zhuyu Peng, Hansruedi Mathys, and Li-Huei Tsai for experimental profiling collaboration and scientific input. The authors additionally thank the NIH NeuroBioBank and the University of Alabama at Birmingham for providing the human HD and control case samples used in this study.

## DATA AVAILABILITY

Count matrices for all 35,757 cells analyzed in this study are uploaded with this submission as Supplementary Data and at http://compbio.mit.edu/scBBB/.

## AUTHOR CONTRIBUTIONS

F.J.G. designed the human and mouse studies and developed the BVE protocol; N.S. conducted data analysis with assistance from F.J.G. and H.L.; B.G. and M.S. assisted in *ex vivo* human tissue sample acquisition; H.L. assisted with BVE snRNA-seq sample preparation; K.G. and J.M. conducted snRNA-seq post-mortem sample profiling; D.A.B. provided *post mortem* samples; F.J.G., N.S., M.K., and M.H. wrote the paper with comments from all authors; and M.K. and M.H. supervised the project.

## Notes

### Competing Interest Statement

The authors have declared no competing interest.

