## Supplemental Figures for "Single-cell dissection of the human cerebrovasculature in health and disease"

**a**

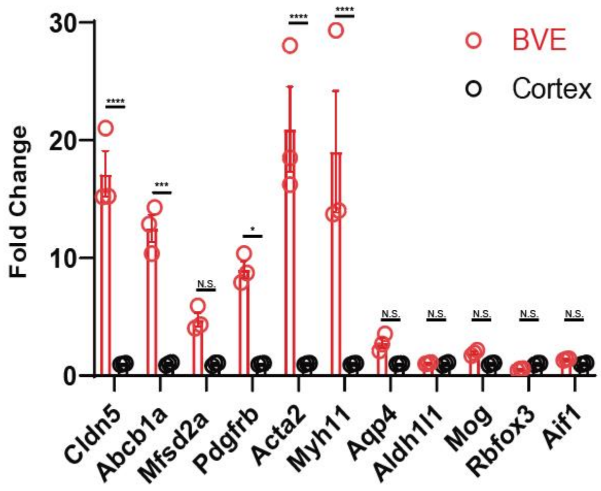


**b**

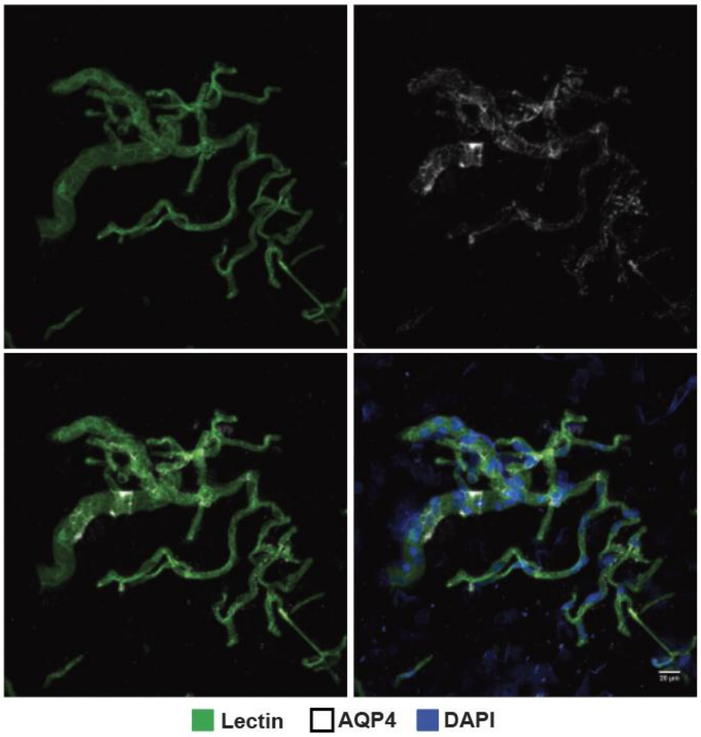


**Extended Data Figure 1. Validation of Blood Vessel Enrichment (BVE) protocol. a.** qPCR of canonical cell type markers for endothelial (*Cldn5, Abcb1a, Mfsd2a*), mural (*Pdgfrb, Acta2, Myh11*), astrocytes (*Aqp4, Aldh1l1*), oligodendrocytes (*Mog*), neurons (*Rbfox3*), and microglia (*Aif1*) from mouse cortex, $n$=3, two-tailed t test, *$p < 0.01$, **$p < 0.001$, ***$p < 0.0001$,****$p < 0.00001$, n.s. = not significant. **b.** Immunofluorescence of blood vessels enriched from mouse cortex using the BVE protocol. Scale bar, 20μm

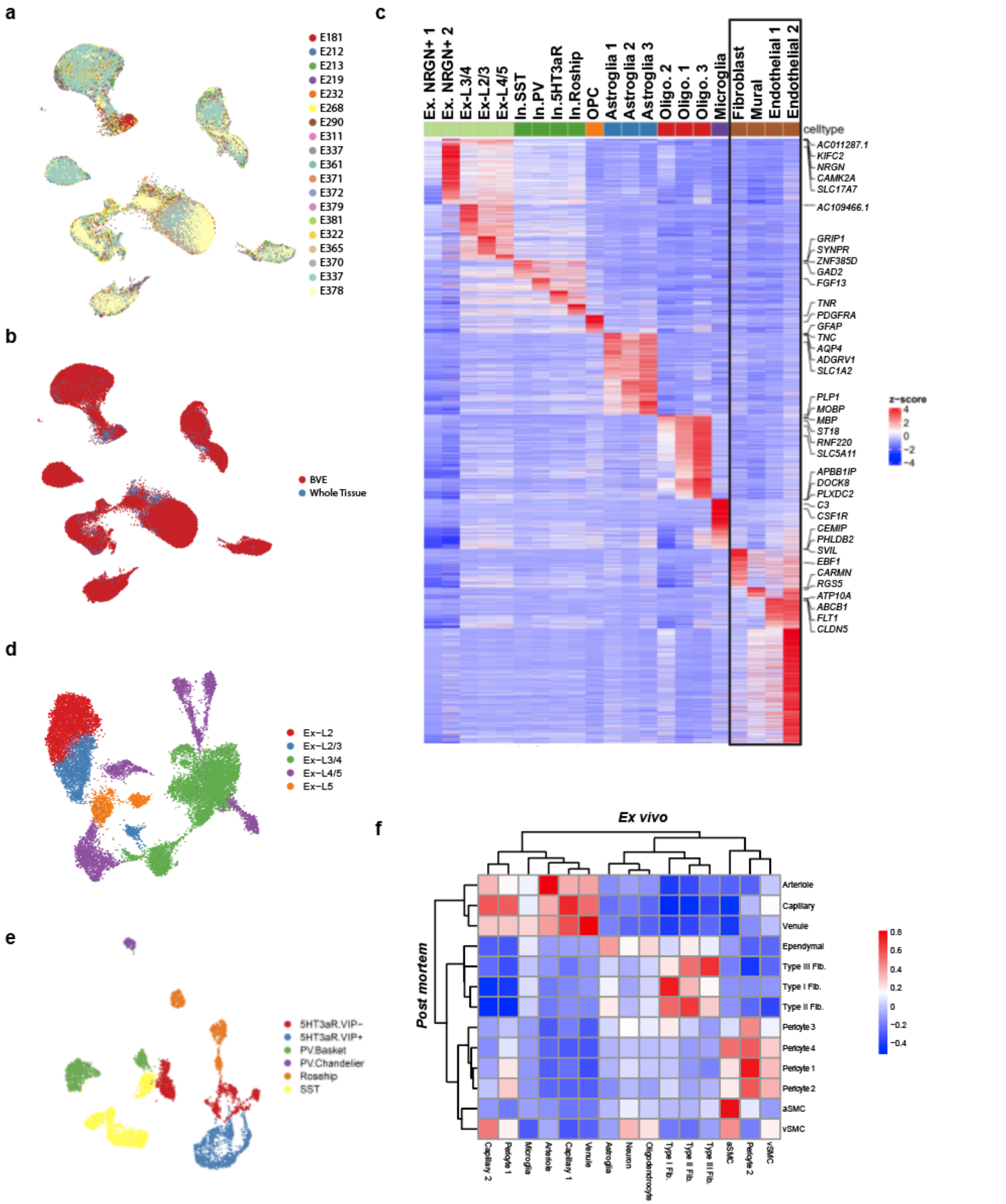


**Extended Data Figure 2. Characterization of human snRNA-seq data from human temporal cortex. a.** UMAP of *ex vivo* dataset by patient ID. **b.** UMAP of *ex vivo* dataset by experimental protocol. **c.** Heatmap of top differentially expressed genes in major cell types from *ex vivo* human tissue. **d.** UMAP sub-clustering of excitatory neurons. **e.** UMAP sub-clustering of inhibitory neurons. **f.** Correlation heatmap between *ex vivo* and *post mortem* vascular cell types

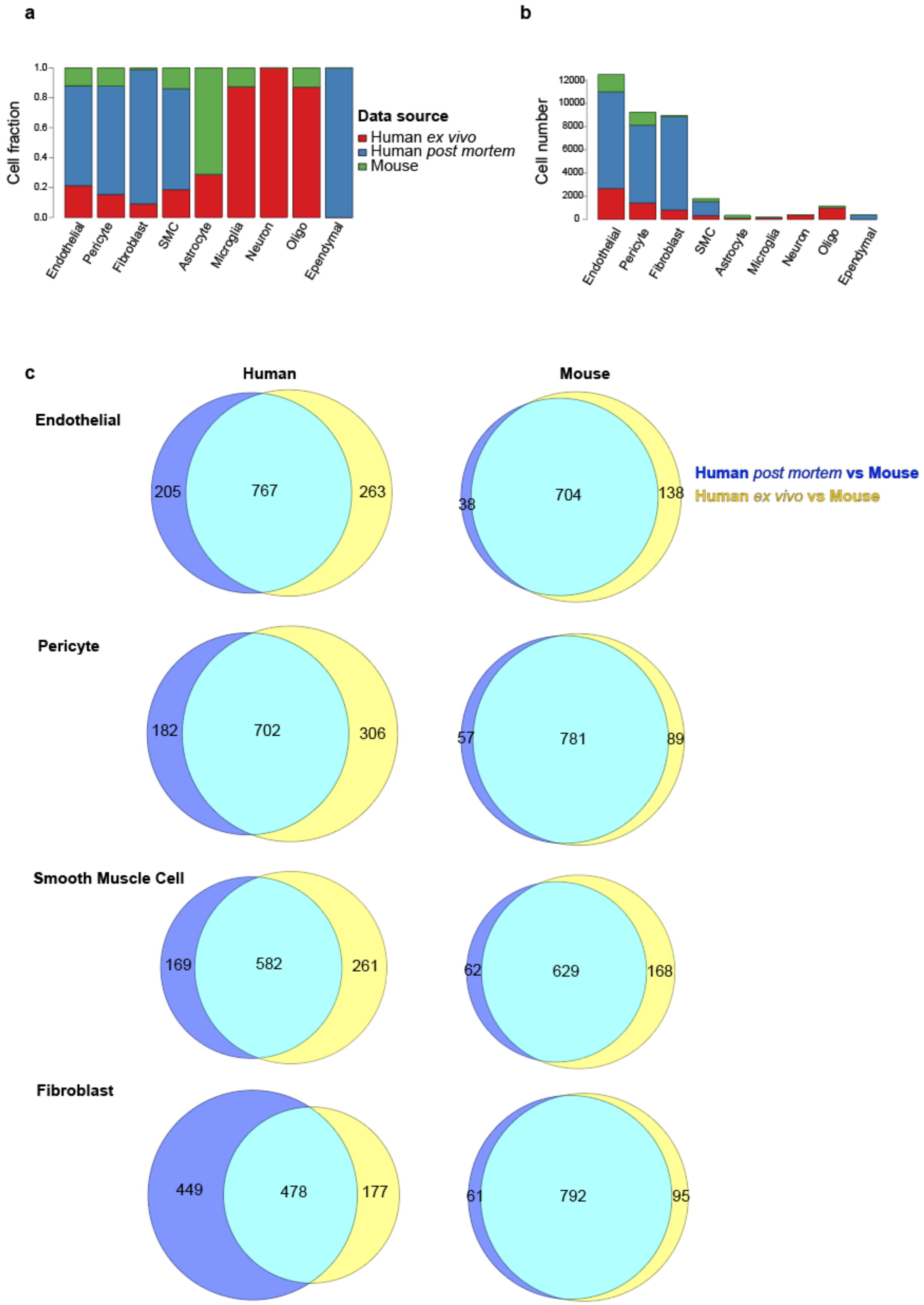


**Extended Data Figure 3. Integrative analysis of *ex vivo*, *post mortem*, and mouse datasets. a.** Cell fraction distribution of single nuclei across all datasets by cerebrovasculature cell type. **b.** Cell number distribution of single nuclei across all datasets by cerebrovasculature cell type. **c.** Venn diagram overlaps of genes between human *post mortem* vs. mouse and human *ex vivo* vs. mouse.

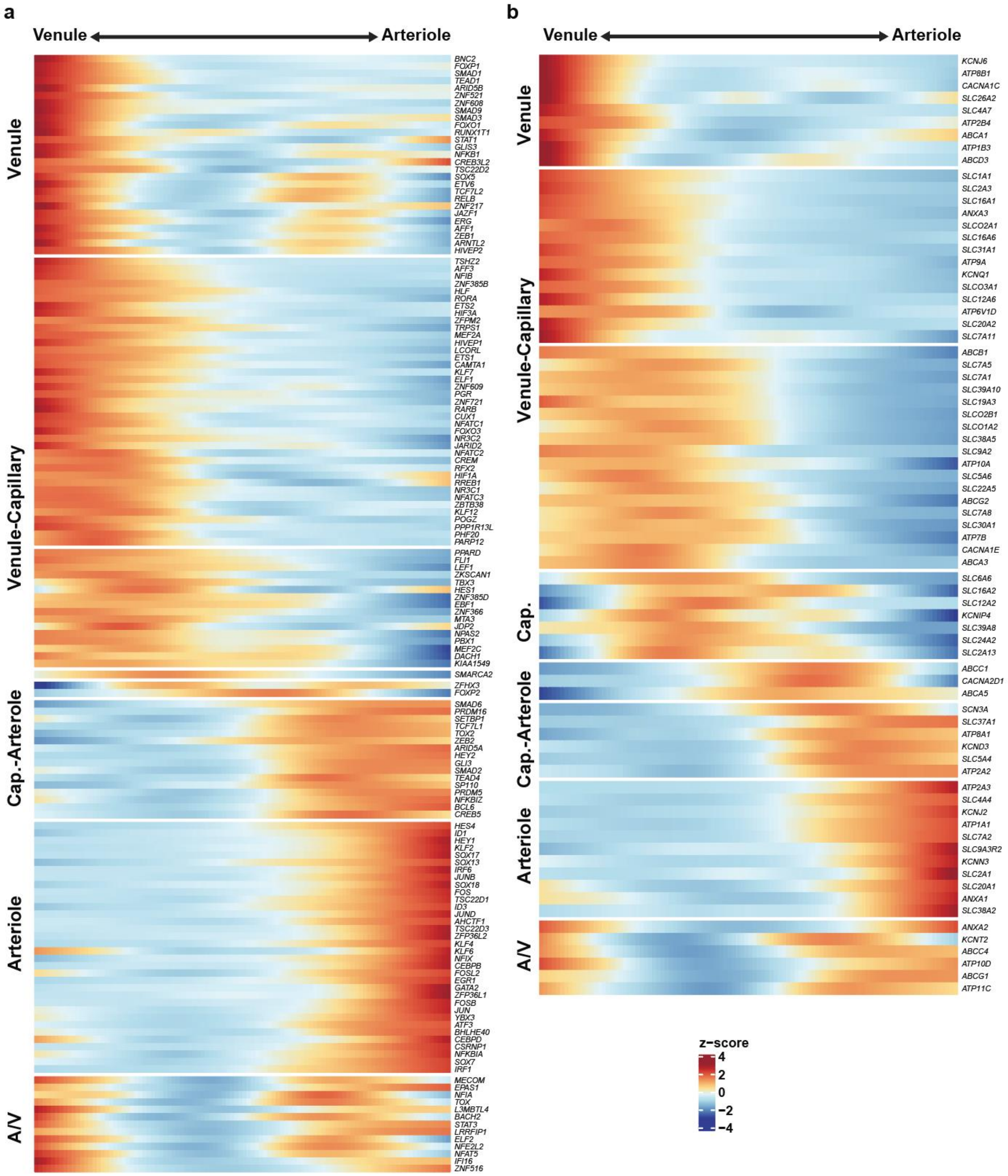


**Extended Data Figure 4. Zonation gene expression analysis of human endothelial cells. a.** Heatmap of 147 zonated transcription factors along the endothelial gradient. **b.** Heatmap of 76 zonated transporters along the endothelial gradient.

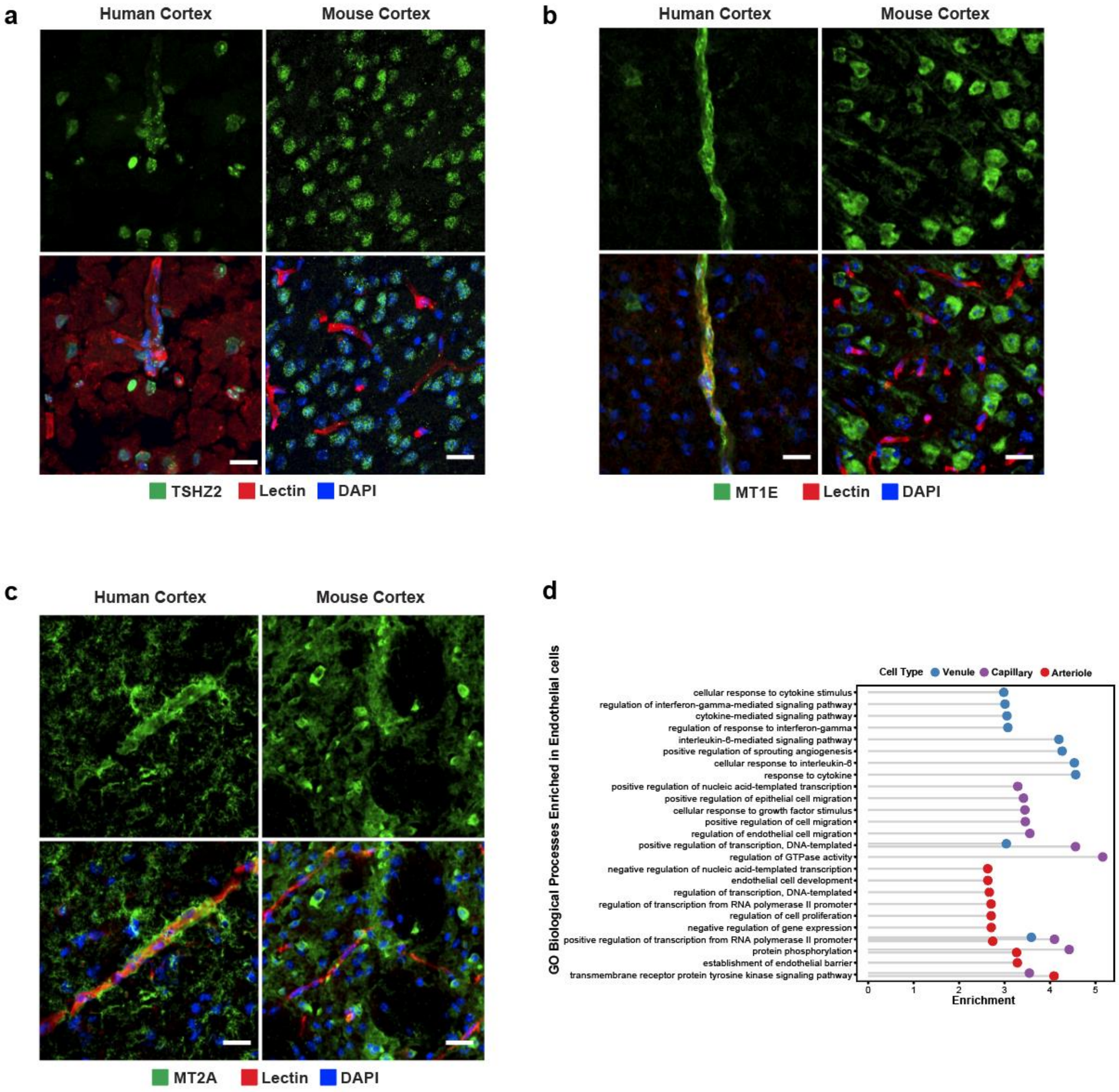


**Extended Data Figure 5. Zonation in human brain endothelial cells. a.** Indirect immunofluorescence of TSHZ2 expression in human and mouse brain cortex. **b.** Indirect immunofluorescence of MT1E/MT1 expression in human and mouse brain cortex. **c.** Indirect immunofluorescence of MT2A/MT2 expression in human and mouse brain cortex. **d.** Enriched Gene Ontology terms in endothelial zones.

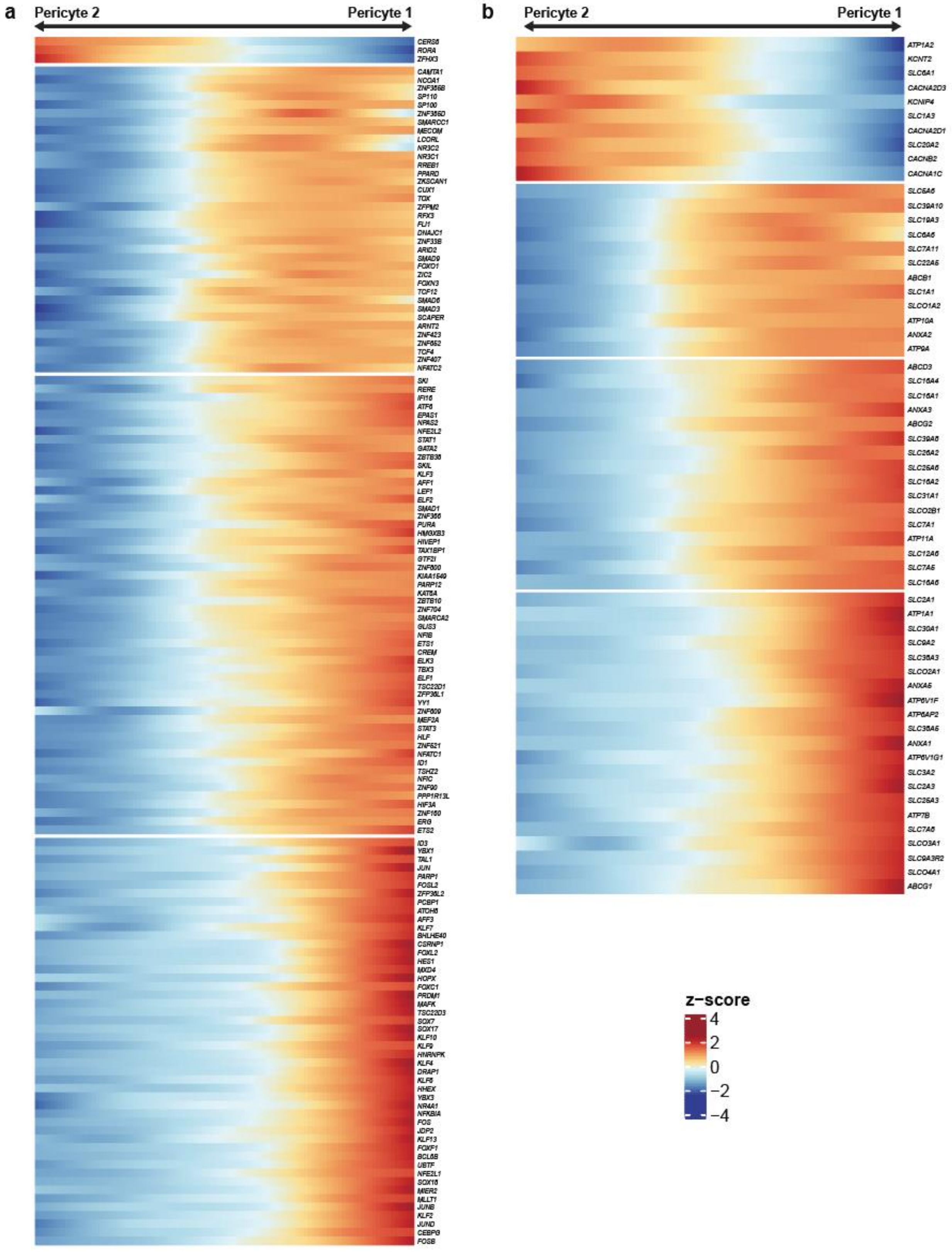


**Extended Data Figure 6. Zonation gene expression analysis of human pericytes. a.** Heatmap of zonated transcription factors along the pericyte gradient. **b.** Heatmap of zonated transporters along the pericyte gradient.

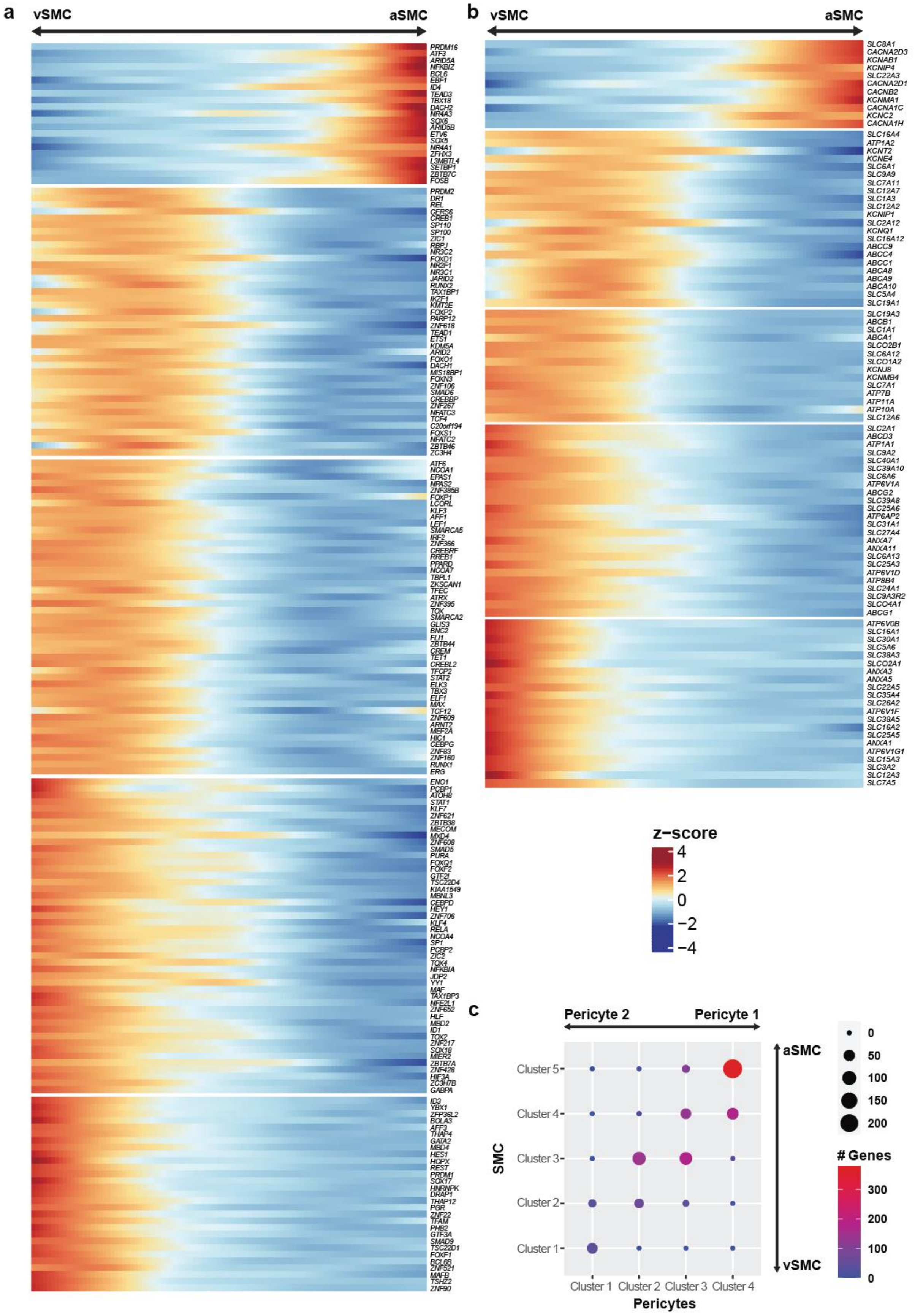


**Extended Data Figure 7. Zonation gene expression analysis of human SMCs. a.** Heatmap of zonated transcription factors along the SMC gradient. **b.** Heatmap of zonated transporters along the SMC gradient. **c.** Overlap matrix across the zonated pericyte and SMC clusters.

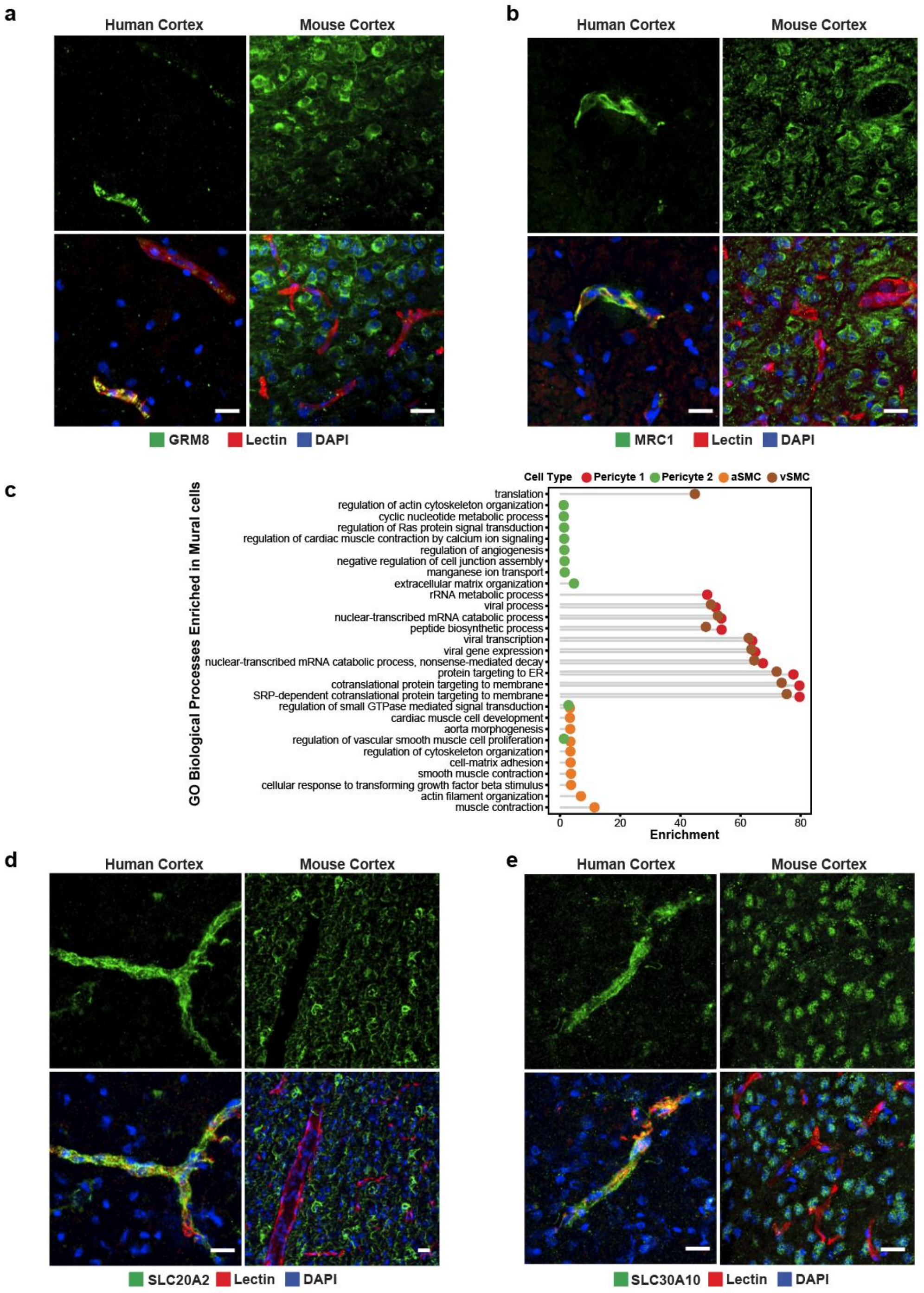


**Extended Data Figure 8. Zonation in human brain mural cells. a.** Indirect immunofluorescence of GRM8 expression in human and mouse brain cortex. **b.** Indirect immunofluorescence of MRC1 expression in human and mouse brain cortex. **c.** Enriched Gene Ontology terms in mural zones. **d.** Indirect immunofluorescence of SLC20A2 expression in human and mouse brain cortex. **e.** Indirect immunofluorescence of SLC30A10 expression in human and mouse brain cortex. Scale bar, 20μm

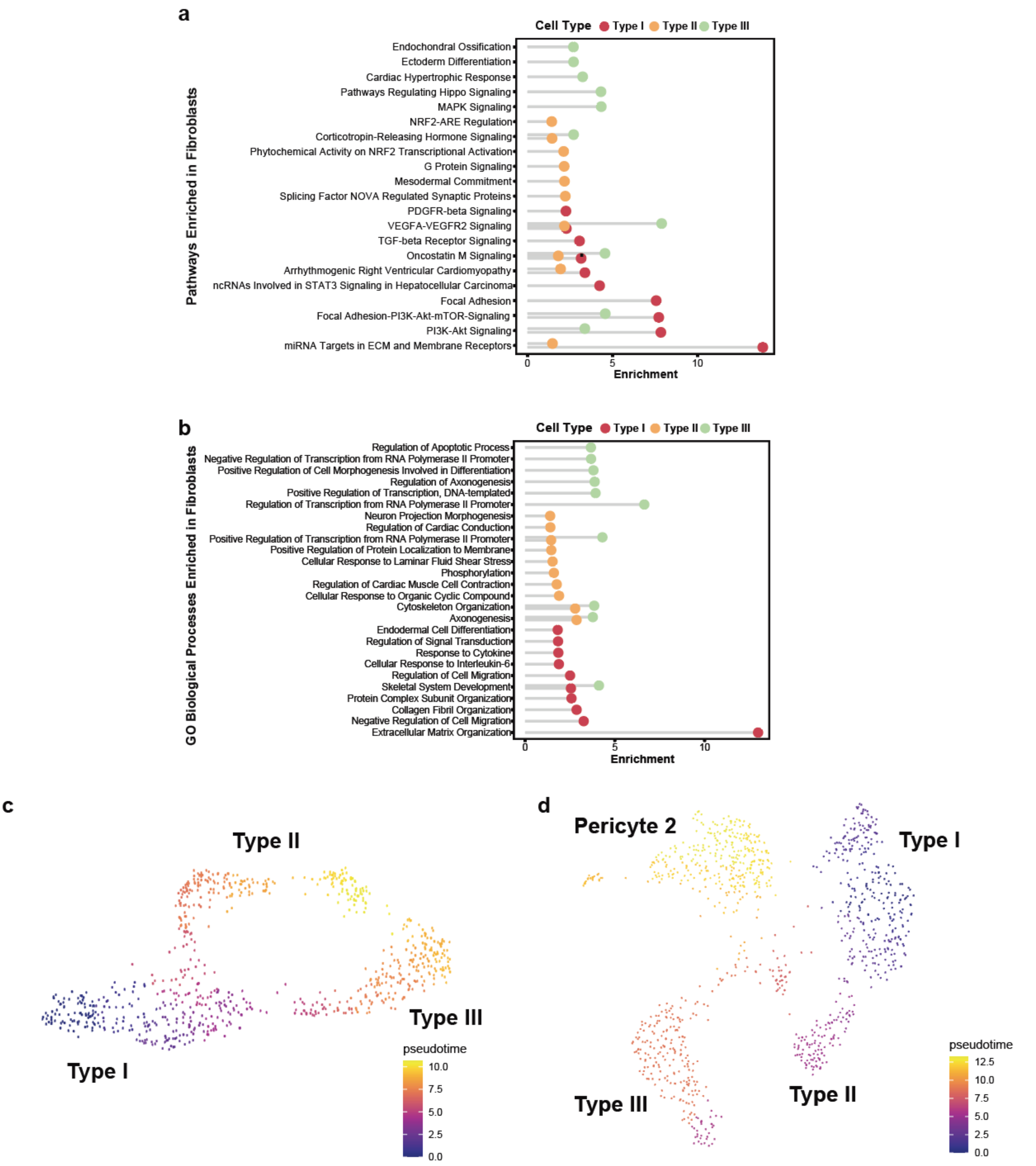


**Extended Data Figure 9. Pathway and Gene Ontology analyses of perivascular fibroblast subtypes. a.** Enriched pathway analysis in perivascular fibroblast subtypes. **b.** Enriched Gene Ontology analysis in perivascular fibroblast subtypes. **c.** Pseudotime analysis of *ex vivo* fibroblast subtypes. **d.** Pseudotime analysis of *ex vivo* fibroblast subtypes and Pericyte 2 (note: Pericyte 1 not shown as it did not fall within any pseudotime trajectory).

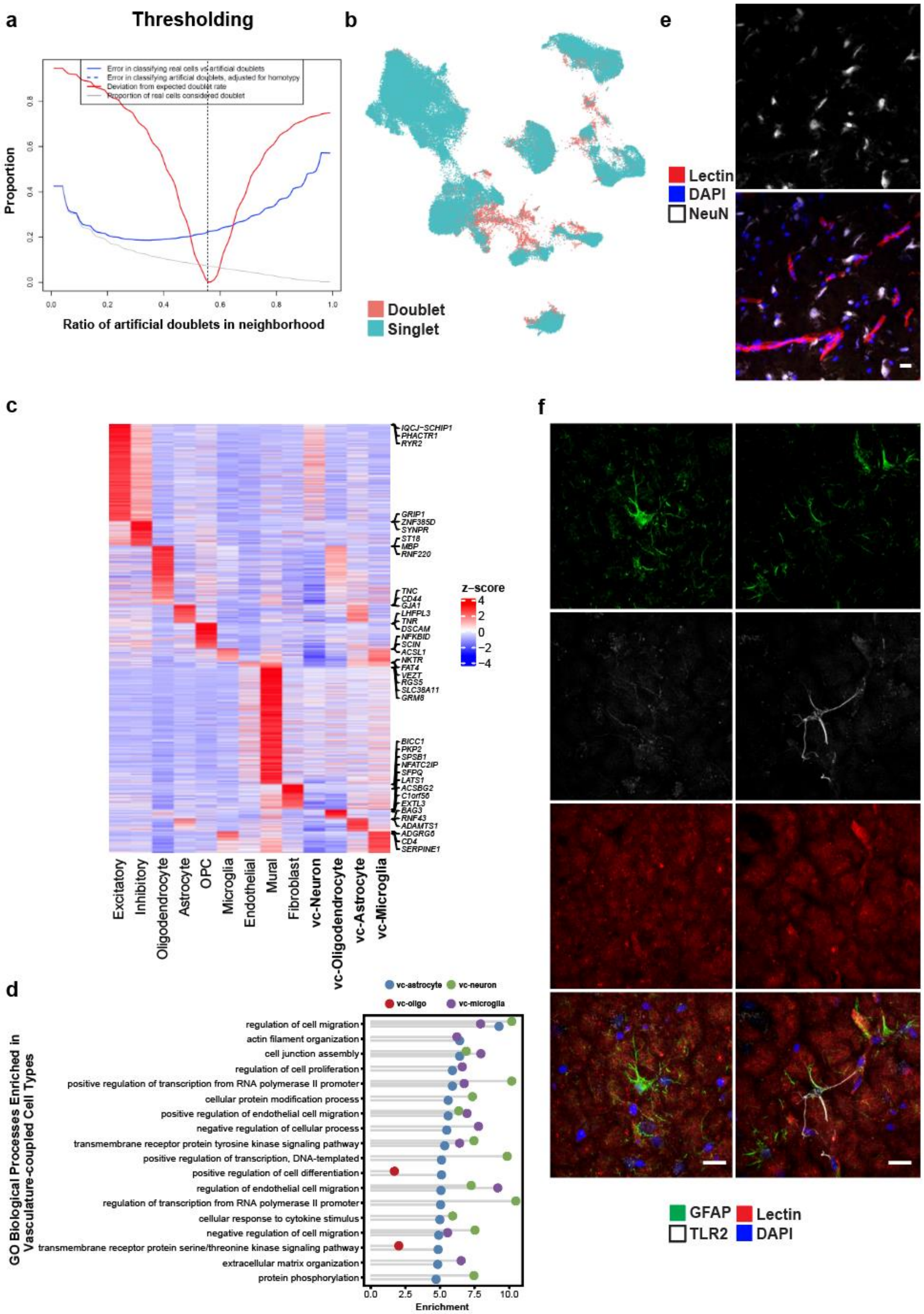


**Extended Data Figure 10. Vascular-coupled (vc) cell types in the *ex vivo* snRNA-seq data. a.** Thresholding parameters for doublet detection. **b.** UMAP of determined singlet/doublets in human *ex vivo* samples. **c.** Heatmap of DEGs in canonical, vascular, and vascular-coupled cell types. **d.** Gene ontology enrichment in vc-cell types. **e.** Immunofluorescence of NeuN (*RBFOX3*)-expressing neurons, with a subset adjacent to blood vessels. **f.** Indirect immunofluorescence staining of *GFAP*-expressing astrocytes (left panels) and *GFAP*-vc-astrocytes (right panels). *TLR2* expressing vc-astrocytes are perivascular compared to canonical astrocytes.

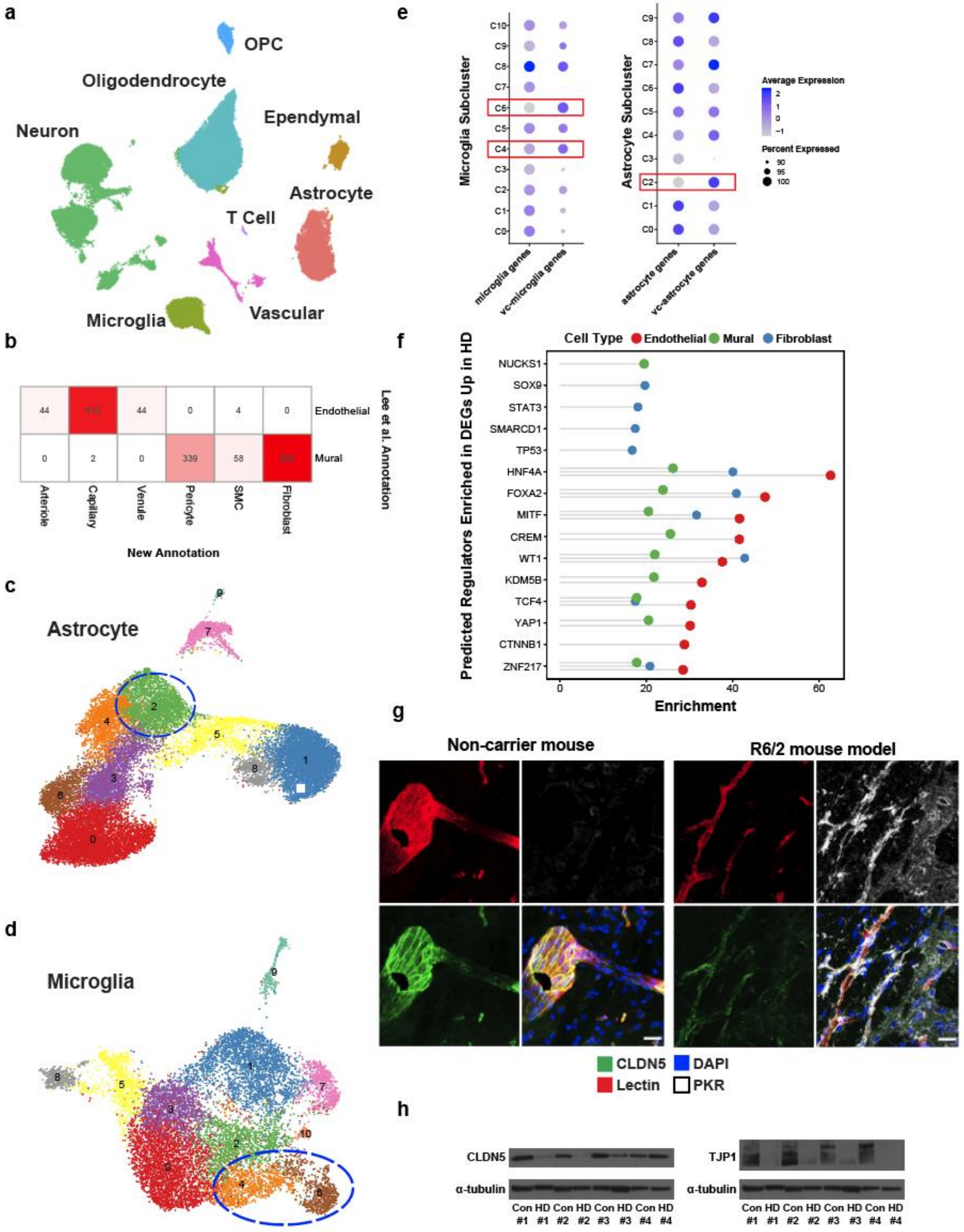


**Extended Data Figure 11. Cerebrovascular profiling in Huntington's disease. a.** UMAP of integrated single nuclei from *post mortem* control and HD human patient samples. **b.** Comparison of cerebrovasculature cell annotations (in cell numbers) in this study vs. Lee et al. **c.** UMAP analysis of astrocyte subclusters in HD. Vasculature-coupled astrocytes outlined in blue. **d.** UMAP analysis of microglia subclusters in HD. Vasculature-coupled microglia outlined in blue. **e.** Dot plot signature score comparison of glial subclusters with respective vc-cell type annotations. Red boxes indicate vc-cell types. **f.** ChEA prediction of top 10 regulators of upregulated genes in HD endothelial, mural, and fibroblasts cells. **g.** PKR immunoreactivity in the R6/2 HD mouse model engulfs blood vessels with low CLDN5 expression. **h.** Western blots for tight junction proteins CLDN5 and TJP1 from human HD and control samples. Scale bar, 20μm.
